# RNA-SeqEZPZ: A Point-and-Click Pipeline for Comprehensive Transcriptomics Analysis with Interactive Visualizations

**DOI:** 10.1101/2024.12.20.629844

**Authors:** Cenny Taslim, Yuan Zhang, Genevieve C. Kendall, Emily R. Theisen

## Abstract

RNA-Seq analysis has become a routine task in numerous genomic research labs, driven by the reduced cost of bulk RNA sequencing experiments. These generate billions of reads that require accurate, efficient, effective, and reproducible analysis. But the time required for comprehensive analysis remains a bottleneck. Many labs rely on in-house scripts, making standardization and reproducibility challenging. To address this, we developed RNA-SeqEZPZ, an automated pipeline with a user-friendly point-and-click interface, enabling rigorous and reproducible RNA-Seq analysis without requiring programming or bioinformatics expertise. For advanced users, the pipeline can also be executed from the command line, allowing customization of steps to suit specific requirements.

This pipeline includes multiple steps from quality control, alignment, filtering, read counting to differential expression and pathway analysis. We offer two different implementations of the pipeline using either (1) bash and SLURM or (2) Nextflow. The two implementation options allow for straightforward installation, making it easy for individuals familiar with either language to modify and/or run the pipeline across various computing environments.

RNA-SeqEZPZ provides an interactive visualization tool using R shiny to easily select the FASTQ files for analysis and compare differentially expressed genes and their functions across experimental conditions. The tools required by the pipeline are packaged into a Singularity image for ease of installation and to ensure replicability. Finally, the pipeline performs a thorough statistical analysis and provides an option to perform batch adjustment to minimize effects of noise due to technical variations across replicates.

RNA-SeqEZPZ is freely available and can be downloaded from https://github.com/cxtaslim/RNA-SeqEZPZ.

## Introduction

Data analysis of RNA-Seq consists of a set of successive stages that are repetitive and routinely executed using a wide variety of tools. Typically, analysis starts with quality control of raw sequence reads, and is followed by alignment of reads to a reference genome, filtering of low-quality reads, counting reads that aligned to a specific feature/gene, differential analysis of genes in different condition and finally visualization of the results^1^. In house analysis usually involves a bioinformatician creating step-by-step scripts for specific datasets which will need to be modified for different datasets. With each modification and customization, it is notoriously challenging to keep the analysis fully reproducible primarily due to differences in scripts, hardware, operating systems, and software versions. Reproducibility is critical in bioinformatics analysis to ensure reliable validation of scientific findings^2^. Furthermore, though many aspects of RNA-seq analysis are routine, bioinformatics analysis remains a significant bottleneck in this process^3^. There is thus demand for an easy-to-run, comprehensive pipeline that expedites routine RNA-Seq analysis without sacrificing the quality and reproducibility of the results. Here, we describe RNA-SeqEZPZ, a point-and-click tool for comprehensive analysis of RNA-seq experiments from raw data to result visualization. The software tools used are packaged and containerized to makes it easier to install and use^4^. RNA-SeqEZPZ is designed to be accessible to a wide range of users, including bench scientists and bioinformaticians. Wet lab scientists will appreciate the user-friendly point-and-click interface, while advanced users can customize the open-source scripts to suit their specific needs.

There are several existing RNA-Seq pipelines, particularly ENCODE^5^ and nf-core^6^ pipelines. In comparison to these pipelines, a notable feature of RNA-SeqEZPZ is its point-and-click interface and interactive visualization capabilities designed to enable comprehensive analysis by bench scientists without formal training in bioinformatics. Several shiny apps providing graphical interface for RNA-Seq analysis such as ROGUE^7^, Shiny-Seq^8^ and bulkAnalyseR^9^ have also been previously published. However, none of these tools support analyzing RNA-Seq experiments starting from raw FASTQ files. We found that Partek™ flow, a commercial software, offers functionalities most similar to RNA-SeqEZPZ. To the best of our knowledge, RNA-SeqEZPZ is the first open-source tool to offer a point-and-click interface with interactive plots, starting from raw FASTQ reads and providing analytical capabilities from differential genes analysis to pathway analysis. This pipeline can potentially accelerate research progress by simplifying a complex process, enhancing reproducibility within and across labs, and empowering researchers to focus on interpreting their own results.

## Results

### Analysis workflow

RNA-SeqEZPZ encompasses multiple steps, utilizes various tools, and produces multiple output files (Figure 1 and Supplementary Figure 1). It accepts gzipped paired-end sequencing FASTQ files as input. Additionally, it supports alignment to reference genomes other than human (i.e. zebrafish and mouse), which can be configured during installation. Users can select all the inputs through a point-and-click interface implemented in a Shiny app^10^ and initiate comprehensive analysis. Specifically, RNA-SeqEZPZ performs the following steps: (1) Merging FASTQ files from different lanes, (2) Creating a genome index if species is not human or mouse, (3) Running quality control, trimming of adapter and low-quality sequences using trim_galore^11^, FASTQC^12^ and MultiQC^13^, (4) Aligning reads using STAR 2-pass approach to improve alignment rate and sensitivity^14^, (5) Creating sequence-depth normalized and replicates-combined bigwig tracks using bamCoverage^15^ and WiggleTools^16^, (6) Counting reads that are assigned to genomic features using featureCounts^17^, and finally, (7) Perform differential gene analysis comparing experiments with different conditions and generating a statistical report of the analysis using DESeq2^18^ and SARTools^19^. A more detailed documentation regarding each step and the installation can be found in https://github.com/cxtaslim/RNA-SeqEZPZ. Below, we describe in more details the components of the RNA-SeqEZPZ pipeline, including interface implementation in Shiny^10^.

**Figure 1.**
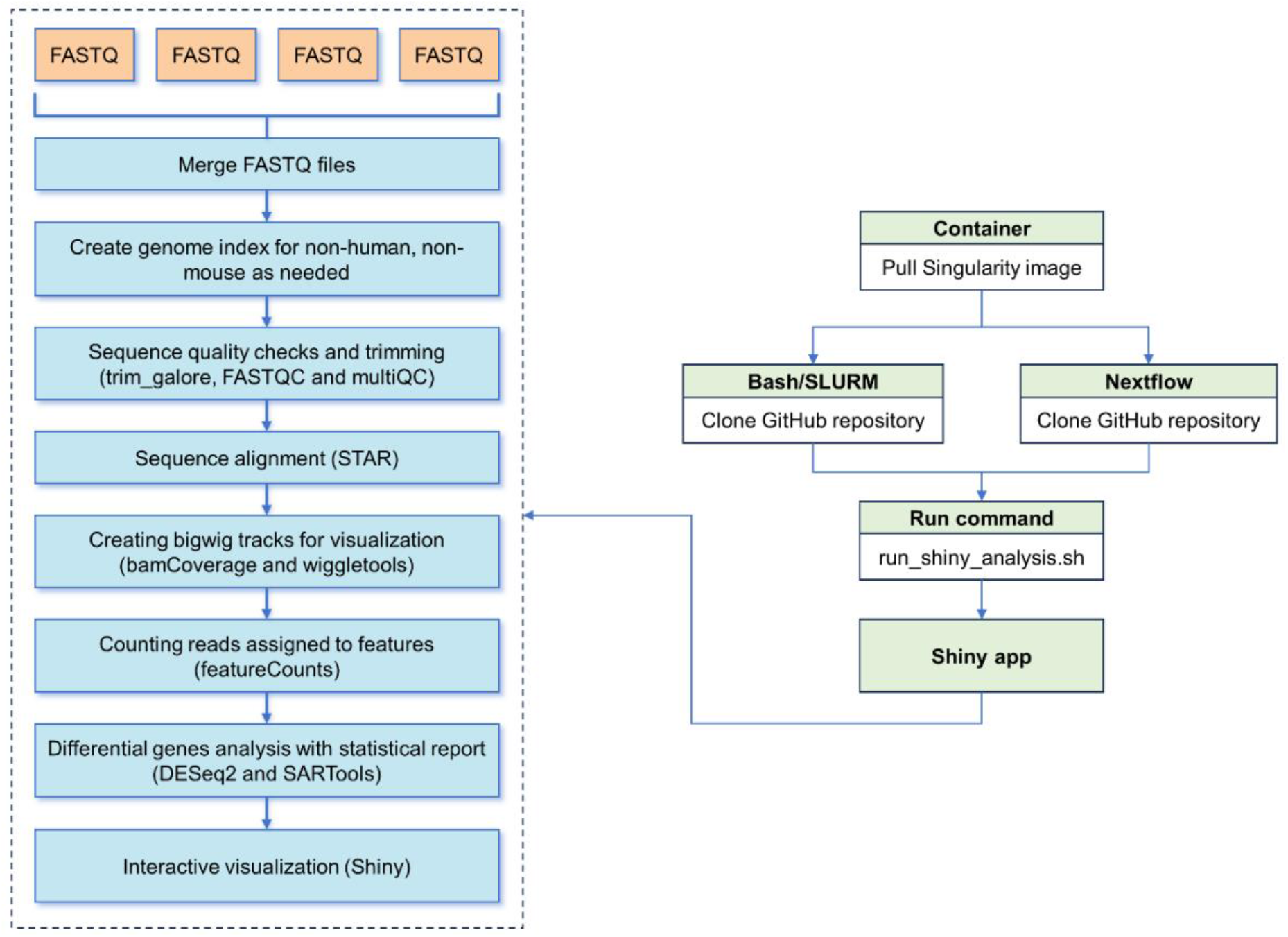
Overview of the analysis steps, tools and the technologies used in RNA-SeqEZPZ

### Software implementation

RNA-SeqEZPZ is a combination of a Shiny^1^ app with either bash scripts and SLURM^20^ (a cluster resource management system) or Nextflow^21^ scripts. It is specifically designed to run on a High-Performance Computing (HPC) system. As SLURM^20^ is the most widely used workload manager in HPC^22^, using it in bash scripts will enable users to easily modify the scripts as needed and leverage their existing familiarity with the system. Nextflow^21^ is a modern workflow management system designed to simplify the development and deployment of complex data analysis pipelines. Nextflow^21^ enhances the flexibility of this pipeline to run on diverse computation infrastructures with workload managers other than SLURM^20^, while also simplifying pipeline implementation. The required R packages, and all other tools needed for analysis are enclosed inside a Singularity^23^ container removing any potential difficulties involved in the installation of all the required software. Altogether this promotes the reproducibility, standardization, and portability of the RNA-seqEZPZ pipeline. Further, the shiny app and analysis run locally on an HPC cluster, removing the need to upload gigabytes to terabytes of data to an outside server over the internet.

### User friendly interface

A primary design goal of RNA-SeqEZPZ is to accelerate full analysis of RNA-seq datasets and provide interactive analysis of the results. As such, the pipeline is designed to be run with a one-line command in the terminal that loads a user-friendly interface implemented as a Shiny^10^ app (Figure 2). The interface is viewed in a Firefox browser, so users can easily zoom in/out, make the text bigger and adjust the size of the window. To run the analysis, users simply select their FASTQ files and provide the necessary information for each file through an intuitive file browser interface. Once all information about the samples has been filled in, clicking on “Run full analysis” will automatically run the full analysis described above (Figure 1).

**Figure 2.**
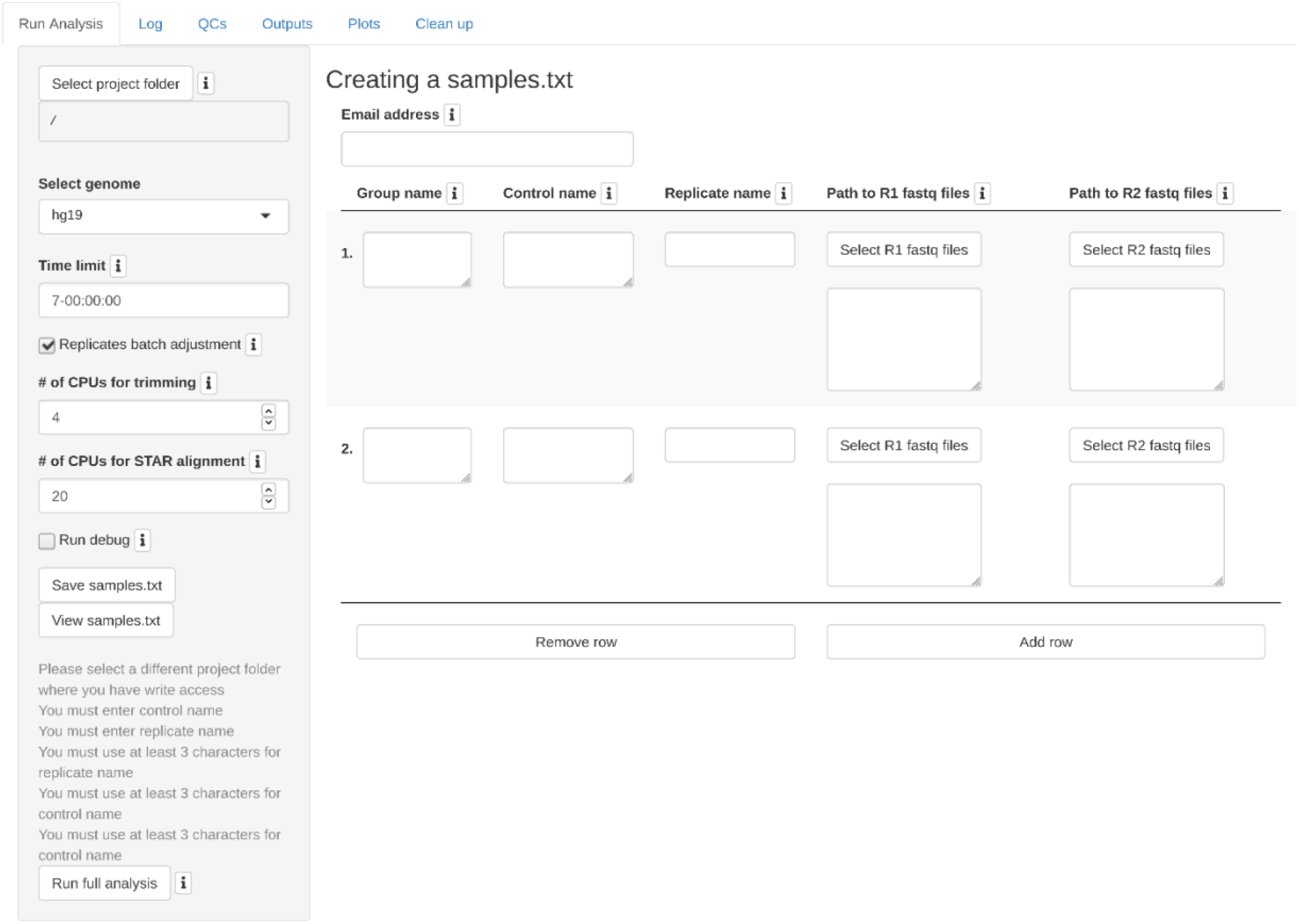
A screenshot of the run analysis interface where users will be able to click-and-select their FASTQ files, reference genome, and other input. There is an “**i**” icon which will provide more information when hovered over in the interface.

While the analysis is ongoing, users will be able to see progress from the “Log” tab (Figure 3).

**Figure 3.**
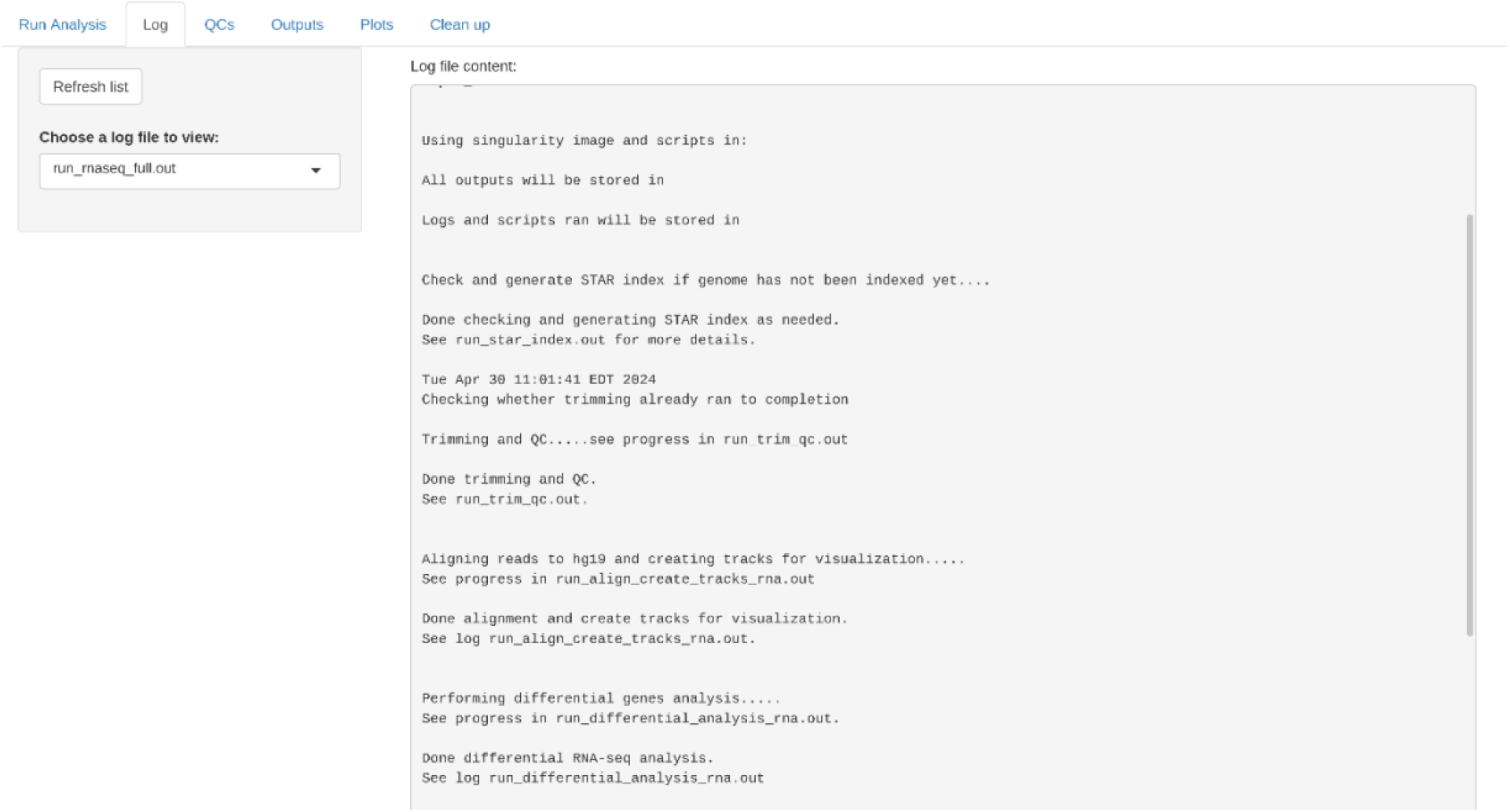
A snapshot of the log file providing information on the current progress of the RNA-Seq analysis.

Once the analysis has completed, users will be able to click on the “QC” tab and see all the quality control metrics compiled by MultiQC^13^ (Figure 4).

**Figure 4.**
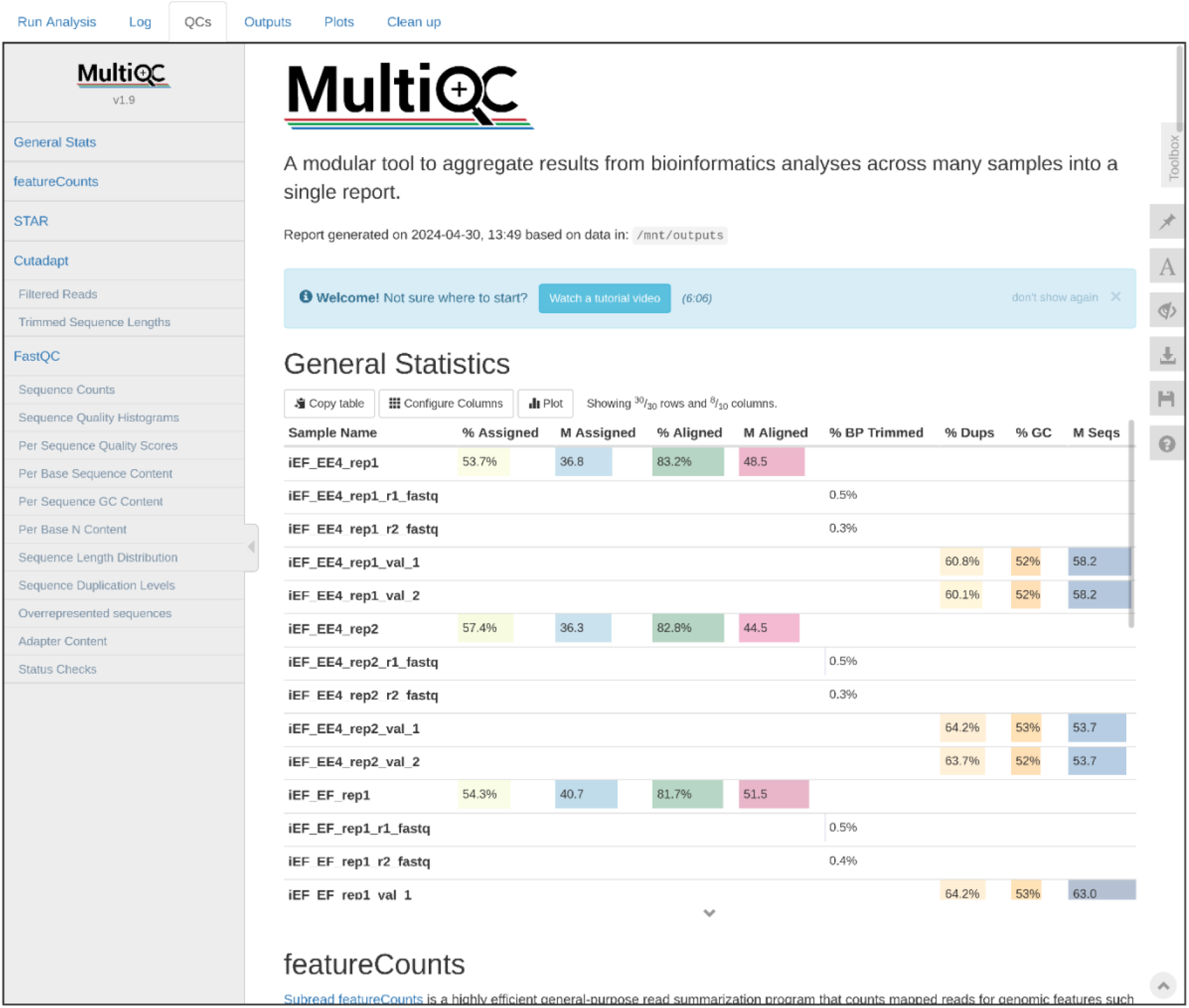
A screenshot of the quality control report in interactive HTML format that can be viewed by users by clicking the “QCs” tab.

The MultiQC^13^ generated HTML files are interactive as well, which allows for some customization of the plots. The QC report includes metrics for raw reads, alignment rate, number of duplicated reads, percentage of reads aligned to genomic features, etc. A statistical report of the differential gene analysis can be viewed in the “Outputs” tab (Figure 5). This report is generated using a modified version of SARTools^19^. The report contains description of raw data, Principal Component Analysis (PCA) plot and hierarchical clustering of samples to explore the variability within and between samples. The statistical report also described the steps performed in the differential analysis using DESeq2^18^ along with the statistical assumptions and validation of the choices used.

**Figure 5.**
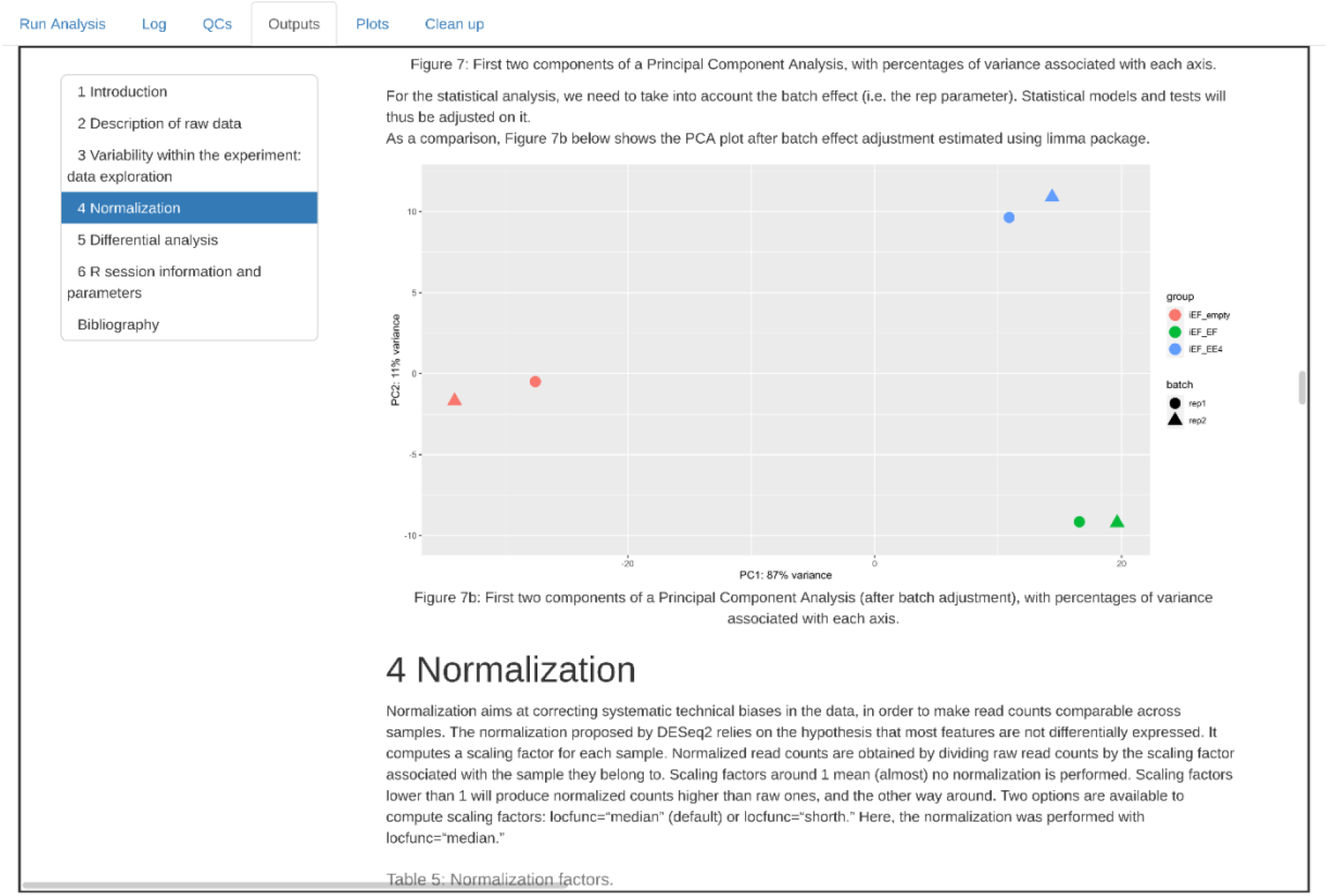
A screenshot of statistical report includes a PCA plot illustrating how the samples cluster across biological replicates and conditions, along with an explanation of the normalization method used on the analysis.

“Plots” tab provides users the ability to adjust the cut-offs for significant differential genes, find the log_2_ fold-change of their gene of interest, create a volcano plot, and perform overlap and pathway analysis (Figure 6 A-E). The GeneOverlap^24^ package was utilized to compute the Jaccard similarity index^25^ and Fisher’s exact test^26^ to evaluate the significance of overlap between the gene lists (not shown). The overlaps between genes in different conditions were visualized using proportional Euler and Venn diagrams, as well as an UpSet plot, created using eulerr^27^, Venn^28^ and UpSetR^29^ packages. Pathway analysis or Over-Representation analysis was conducted using clusterProfiler^30^ and msigdbr^31^ packages. All other plots were generated using ggplot2^32^ package.

**Figure 6.**
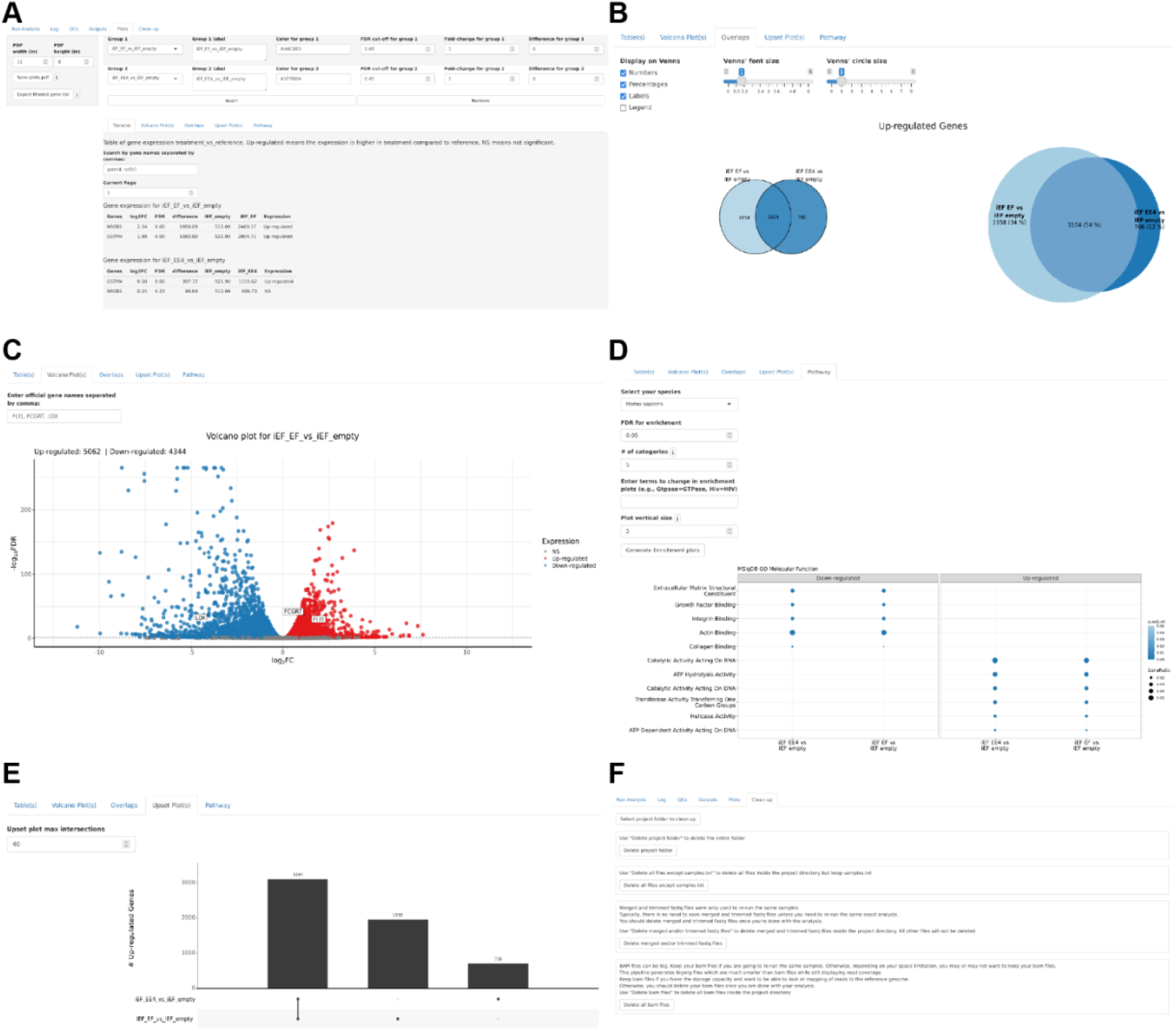
Tables and interactive plots. (A) Table showing the sorted log_2_ Fold-Change, False Discovery Rate (FDR) and read counts difference between treatment and control samples. (B) Overlap of up-regulated genes in two or more groups. (C) Volcano plot with user highlighted gene of interest. (D) Pathway analysis using msigdb databases. (E) Upset plot showing the overlap of up-regulated genes. (F) A cleanup interface allows users to delete large files such as merged and trimmed FASTQ and BAM files.

Additionally, since the files, including intermediate ones generated by the pipeline can accumulate to terabytes in size, we provide users a simple way to delete projects and files they no longer require (Figure 6F). We also provide explanations to help users determine whether to keep or delete these files.

### Installation and usage

Installation instructions are provided in detail at https://github.com/cxtaslim/RNA-SeqEZPZ. Briefly, a Singularity image is pulled from a repository. Then, depending on user preference, either a Nextflow or a bash/SLURM repository containing all the necessary scripts should be cloned (Figure 1). To use the pipeline, users need to connect to their HPC cluster and run a one-line command: “`run_shiny_analysis.sh`” which will bring up a Firefox browser where user will be able to select the sample FASTQ file path, output path, resource requirements, and various settings. Options are also available for running the steps of the pipeline individually (see the manual on the website for details). To assist users running this for the first time, we have provided example datasets that can be downloaded from https://github.com/cxtaslim/RNA-SeqEZPZ, along with an easy-to-follow step-by-step tutorial.

### Public dataset analysis

In order to show the utility of the pipeline, we analyzed the RNA-Seq experiments in the study of novel Ewing Sarcoma fusion proteins^33^. RNA-sequencing analysis was performed on two biological replicates from a knockdown/rescue experiment in the A673 human cell line where the endogenous fusion oncogenic transcription factor EWSR1::FLI1 was depleted by shRNA (iEF) and then rescued with either EWSR1::FLI1 (iEF + EF) or EWSR1::ETV4 (iEF + EE4) constructs. These samples were compared to control cells with no rescue (iEF + Empty Vector). The FASTQ files can be downloaded from GEO (GSE173185). As shown in the QC report, there are 48.5 million reads that are aligned with 83.4% alignment rate and 53.7% of the reads are assigned to a feature (Figure 4 and Supplementary File 1). The statistical report shows the PCA plot where the 6 samples are clustered by replicates and rescue condition (Figure 5 and Supplementary File 2). Figure 6A shows the shiny interface comparing the different rescue conditions, with a screenshot of the two well-known target genes (i.e. NR0B1 and GSTM4) in cell line rescued with EWSR1::FLI1 and EWSR1::ETV4 constructs. The Euler diagram shows significant overlap of 3,104 genes that are both up-regulated by EWSR1::FLI1 and EWSR1::ETV4 (Figure 6B and Supplementary File 3). The volcano plot highlights FLI1 (indicative of rescued EWSR1::FLI1 expression), and FCGRT and LOX which are known to be up- and down-regulated by EWSR1::FLI1^34,35^, respectively (Figure 6C and Supplementary File 3). Volcano plot for genes regulated by EWSR1::ETV4 is also provided (not shown). Pathway analysis indicates that genes regulated by EWSR1::ETV4 and EWSR1::FLI1 are involved in many similar functions (Figure 6D and Supplementary File 3). EWSR1::FLI1 downregulated genes are consistent with those identified in a previous study by Kinsey *et al*.^36^ (Supplementary File 3). The QC report (Supplementary File 1) and statistical report of the differential analysis (Supplementary File 2) are saved as HTML files. All the plots created in RNA-SeqEZPZ can be exported as a pdf file (Supplementary File 3). One of the most commonly used outputs is the list of differentially expressed genes. These tables list genes that are defined as significant along with their Ensembl ID, raw and normalized read count, fold-changes, p-values adjusted for multiple testing, and other statistics generated by the DESeq2 models (Supplementary File 4). Video tutorial on the analysis of this dataset is included in Supplementary File 5.

## Discussions

In summary, RNA-SeqEZPZ provides an easy point-and-click comprehensive analysis of RNA-Seq data which enables biologists to analyze and explore the nuances of their own experiments. The implementation of RNA-SeqEZPZ ensures reproducible analysis and is broadly flexible for running in various computational infrastructures. RNA-SeqEZPZ also provides an entry point analysis for more advanced users where they can download the results and do additional downstream analysis or modify the pipeline to include more features. Thus, RNA-SeqEZPZ represents a valuable easy-to-use tool for the scientific community, enabling the analysis, interpretation, and discovery of insights about gene function and regulation through RNA-Seq experiments.

### Additional files

Supplementary File 1: QC report as generated by RNA-SeqEZPZ in interactive html format. Link to Supplementary Files

Supplementary File 2: Statistical report of the differential gene analysis as generated by RNA-SeqEZPZ.

Supplementary File 3: Plots in pdf format as created in RNA-SeqEZPZ. Link to Supplementary Files

Supplementary File 4: An example of a list of up-regulated genes comparing knockdown/rescue with EWSR1::FLI1 construct to the empty construct as generated by RNA-SeqEZPZ.

Supplementary File 5: Video tutorial of RNA-SeqEZPZ. Link to Supplementary Files

Supplementary Figure 1: RNA-SeqEZPZ workflow showing output files generated. Some icons were sourced and/or adapted from https://nf-co.re/dualrnaseq, created by Regan Hayward under the MIT license.

***Supplementary Figure 1***: *RNA-SeqEZPZ workflow with output files generated*.

## Data Availability

No new data were generated for this study. The data used in this article are available in NCBI Gene Expression Omnibus (GEO) repository (https://www.ncbi.nlm.nih.gov/geo/), under the accession number GSE173185.

## Acknowledgements

This research was partially supported by the High Performance Computing Facility at the Abigail Wexner Research Institute, Nationwide Children’s Hospital.

## Author Contributions

C.T, Y.Z., G.C.K, E.R.T. conceived the main idea, the framework of the pipeline and the manuscript. C.T, Y.Z., G.C.K, E.R.T drafted and improved the manuscript. C.T and Y.Z. developed and implemented the pipeline. G.C.K and E.R.T revised the manuscript, supervised the development of the pipeline, and provided funding. All authors read and commented on the manuscript.

