## Supplementary figures and images for "RNA-SeqEZPZ: A Point-and-Click Pipeline for Comprehensive Transcriptomics Analysis with Interactive Visualizations"

### Supplementary Figure 1

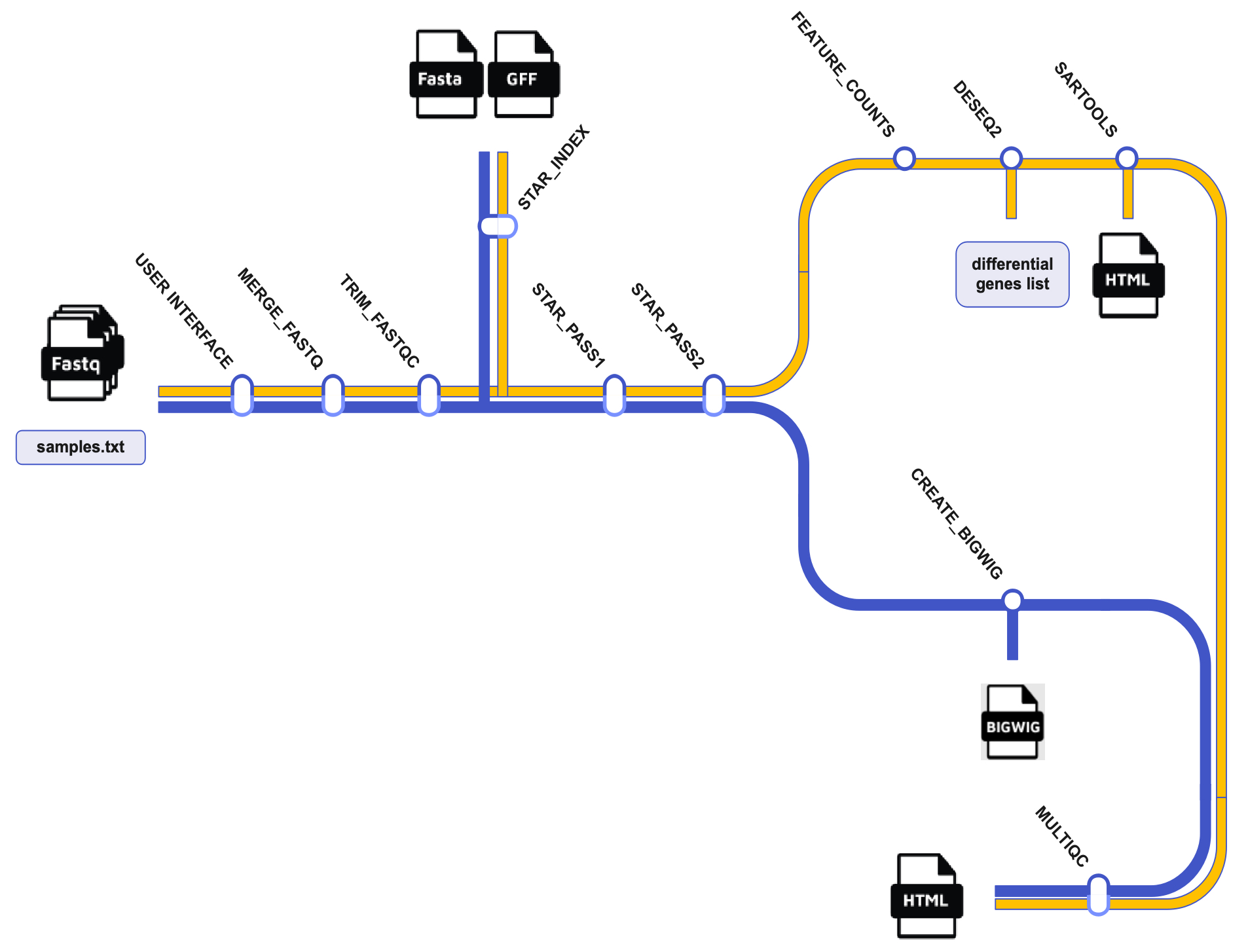
