## Supplementary File 2 for "RNA-SeqEZPZ: A Point-and-Click Pipeline for Comprehensive Transcriptomics Analysis with Interactive Visualizations": Supplementary File 2.html

Statistical report of project RNA-seq\_differential\_analysis: pairwise comparison(s) of conditions with DESeq2


### Statistical report of project RNA-seq\_differential\_analysis: pairwise comparison(s) of conditions with DESeq2

#### NA

#### 2024-04-30

The SARTools R package which generated this report has been developped at PF2 - Institut Pasteur by M.-A. Dillies and H. Varet and modified by Cenny Taslim. Thanks to cite H. Varet, L. Brillet-Guéguen, J.-Y. Coppee and M.-A. Dillies, *SARTools: A DESeq2- and EdgeR-Based R Pipeline for Comprehensive Differential Analysis of RNA-Seq Data*, PLoS One, 2016, doi: http://dx.doi.org/10.1371/journal.pone.0157022 when using this tool for any analysis published.

### 1 Introduction

The analyses reported in this document are part of the RNA-seq\_differential\_analysis project. The aim is to find features that are differentially expressed between iEF\_empty, iEF\_EF and iEF\_EE4. The statistical analysis process includes data normalization, graphical exploration of raw and normalized data, test for differential expression for each feature between the conditions, raw p-value adjustment and export of lists of features having a significant differential expression between the conditions. In this analysis, the rep effect will be taken into account in the statistical models.

The analysis is performed using the R software [1], Bioconductor [2] packages including DESeq2 [3,4] and the SARTools package developed at PF2 - Institut Pasteur. Normalization and differential analysis are carried out according to the DESeq2 model and package. This report comes with additional tab-delimited text files that contain lists of differentially expressed features.

For more details about the DESeq2 methodology, please refer to its related publications [3,4].

### 2 Description of raw data

The count data files and associated biological conditions are listed in the following table.

Table 1: Data files and associated biological conditions.

| label | files | group | rep |
| --- | --- | --- | --- |
| iEF\_emptyrep1 | iEF\_empty\_rep1\_counts.txt | iEF\_empty | rep1 |
| iEF\_emptyrep2 | iEF\_empty\_rep2\_counts.txt | iEF\_empty | rep2 |
| iEF\_EFrep1 | iEF\_EF\_rep1\_counts.txt | iEF\_EF | rep1 |
| iEF\_EFrep2 | iEF\_EF\_rep2\_counts.txt | iEF\_EF | rep2 |
| iEF\_EE4rep1 | iEF\_EE4\_rep1\_counts.txt | iEF\_EE4 | rep1 |
| iEF\_EE4rep2 | iEF\_EE4\_rep2\_counts.txt | iEF\_EE4 | rep2 |

After loading the data we first have a look at the raw data table itself. The data table contains one row per annotated feature and one column per sequenced sample. Row names of this table are feature IDs (unique identifiers). The table contains raw count values representing the number of reads that map onto the features. For this project, there are 63677 features in the count data table.

Table 2: Partial view of the count data table.

|  | iEF\_emptyrep1 | iEF\_emptyrep2 | iEF\_EFrep1 | iEF\_EFrep2 | iEF\_EE4rep1 | iEF\_EE4rep2 |
| --- | --- | --- | --- | --- | --- | --- |
| ENSG00000000003 | 1707 | 2288 | 1725 | 2546 | 1786 | 2481 |
| ENSG00000000005 | 8 | 18 | 8 | 9 | 3 | 1 |
| ENSG00000000419 | 1769 | 1214 | 2055 | 2268 | 1981 | 1657 |
| ENSG00000000457 | 407 | 502 | 398 | 483 | 366 | 406 |
| ENSG00000000460 | 810 | 542 | 1419 | 1301 | 1075 | 865 |
| ENSG00000000938 | 0 | 1 | 1 | 4 | 3 | 3 |

Looking at the summary of the count table provides a basic description of these raw counts (min and max values, median, etc).

Table 3: Summary of the raw counts.

|  | Min. | 1st Qu. | Median | Mean | 3rd Qu. | Max. |
| --- | --- | --- | --- | --- | --- | --- |
| iEF\_emptyrep1 | 0 | 0 | 1 | 576 | 22 | 1210491 |
| iEF\_emptyrep2 | 0 | 0 | 0 | 537 | 19 | 1693581 |
| iEF\_EFrep1 | 0 | 0 | 0 | 639 | 20 | 765960 |
| iEF\_EFrep2 | 0 | 0 | 0 | 681 | 18 | 962760 |
| iEF\_EE4rep1 | 0 | 0 | 1 | 578 | 22 | 704011 |
| iEF\_EE4rep2 | 0 | 0 | 0 | 571 | 18 | 844050 |

Figure 1 shows the total number of mapped and counted reads for each sample. We expect total read counts to be similar within conditions, they may be different across conditions. Total counts sometimes vary widely between replicates. This may happen for several reasons, including:

- different rRNA contamination levels between samples (even between biological replicates);
- slight differences between library concentrations, since they may be difficult to measure with high precision.

Figure 1: Number of mapped reads per sample. Colors refer to the biological condition of the sample.

Figure 2 shows the percentage of features with no read count in each sample. We expect this percentage to be similar within conditions. Features with null read counts in the 6 samples are left in the data but are not taken into account for the analysis with DESeq2. Here, 24265 features (38.11%) are in this situation (dashed line). Results for those features (fold-change and p-values) are set to NA in the results files.

Figure 2: Percentage of features with null read counts in each sample.

Figure 3 shows the distribution of read counts for each sample (on a log scale to improve readability). Again we expect replicates to have similar distributions. In addition, this figure shows if read counts are preferably low, medium or high. This depends on the organisms as well as the biological conditions under consideration.

Figure 3: Density distribution of read counts.

It may happen that one or a few features capture a high proportion of reads (up to 20% or more). This phenomenon should not influence the normalization process. The DESeq2 normalization has proved to be robust to this situation [Dillies, 2012]. Anyway, we expect these high count features to be the same across replicates. They are not necessarily the same across conditions. Figure 4 and table 4 illustrate the possible presence of such high count features in the data set. Please check to make sure the feature with high proportion of reads (usually 20% or more) is NOT rRNA. If it is, you may have contamination.

Figure 4: Percentage of reads associated with the sequence having the highest count (provided in each box on the graph) for each sample.

Table 4: Percentage of reads associated with the sequences having the highest counts.

|  | ENSG00000115414 | ENSG00000258486 | ENSG00000202198 | ENSG00000259001 |
| --- | --- | --- | --- | --- |
| iEF\_emptyrep1 | 3.30 | 1.89 | 1.56 | 1.12 |
| iEF\_emptyrep2 | 4.95 | 1.71 | 1.96 | 0.95 |
| iEF\_EFrep1 | 0.67 | 1.88 | 1.66 | 1.00 |
| iEF\_EFrep2 | 0.83 | 2.22 | 2.15 | 1.41 |
| iEF\_EE4rep1 | 0.88 | 1.91 | 1.65 | 1.01 |
| iEF\_EE4rep2 | 1.04 | 2.19 | 2.32 | 1.34 |

We may wish to assess the similarity between samples across conditions. A pairwise scatter plot is produced (figure 5) to show how replicates and samples from different biological conditions are similar or different (using a log scale). Moreover, as the Pearson correlation has been shown not to be relevant to measure the similarity between replicates, the SERE statistic has been proposed as a similarity index between RNA-Seq samples [5]. It measures whether the variability between samples is random Poisson variability or higher. Pairwise SERE values are printed in the lower triangle of the pairwise scatter plot. The value of the SERE statistic is:

- 0 when samples are identical (no variability at all: this may happen in the case of a sample duplication);
- 1 for technical replicates (technical variability follows a Poisson distribution);
- greater than 1 for biological replicates and samples from different biological conditions (biological variability is higher than technical one, data are over-dispersed with respect to Poisson). The higher the SERE value, the lower the similarity. It is expected to be lower between biological replicates than between samples of different biological conditions. Hence, the SERE statistic can be used to detect inversions between samples.

Figure 5: Pairwise comparison of samples (not produced when more than 12 samples).

### 3 Variability within the experiment: data exploration

The main variability within the experiment is expected to come from biological differences between the samples. This can be checked in two ways. The first one is to perform a hierarchical clustering of the whole sample set. This is performed after a transformation of the count data which can be either a Variance Stabilizing Transformation (VST) or a regularized log transformation (rlog) [3,4].

A VST is a transformation of the data that makes them homoscedastic, meaning that the variance is then independent of the mean. It is performed in two steps: (i) a mean-variance relationship is estimated from the data with the same function that is used to normalize count data and (ii) from this relationship, a transformation of the data is performed in order to get a dataset in which the variance is independent of the mean. The homoscedasticity is a prerequisite for the use of some data analysis methods, such as hierarchical clustering or Principal Component Analysis (PCA). The regularized log transformation is based on a GLM (Generalized Linear Model) on the counts and has the same goal as a VST but is more robust in the case when the size factors vary widely.

Figure 6 shows the dendrogram obtained from VST-transformed data. An euclidean distance is computed between samples, and the dendrogram is built upon the Ward criterion. We expect this dendrogram to group replicates and separate biological conditions.


Figure 6: Sample clustering based on normalized data.

Another way of visualizing the experiment variability is to look at the first principal components of the PCA, as shown on the figure 7. On this figure, the first principal component (PC1) is expected to separate samples from the different biological conditions, meaning that the biological variability is the main source of variance in the data.


Figure 7: First two components of a Principal Component Analysis, with percentages of variance associated with each axis.

For the statistical analysis, we need to take into account the batch effect (i.e. the rep parameter). Statistical models and tests will thus be adjusted on it.  
As a comparison, Figure 7b below shows the PCA plot after batch effect adjustment estimated using limma package.


Figure 7b: First two components of a Principal Component Analysis (after batch adjustment), with percentages of variance associated with each axis.

### 4 Normalization

Normalization aims at correcting systematic technical biases in the data, in order to make read counts comparable across samples. The normalization proposed by DESeq2 relies on the hypothesis that most features are not differentially expressed. It computes a scaling factor for each sample. Normalized read counts are obtained by dividing raw read counts by the scaling factor associated with the sample they belong to. Scaling factors around 1 mean (almost) no normalization is performed. Scaling factors lower than 1 will produce normalized counts higher than raw ones, and the other way around. Two options are available to compute scaling factors: locfunc=“median” (default) or locfunc=“shorth.” Here, the normalization was performed with locfunc=“median.”

Table 5: Normalization factors.

|  | iEF\_emptyrep1 | iEF\_emptyrep2 | iEF\_EFrep1 | iEF\_EFrep2 | iEF\_EE4rep1 | iEF\_EE4rep2 |
| --- | --- | --- | --- | --- | --- | --- |
| Size factor | 0.97 | 0.86 | 1.09 | 1.13 | 1.05 | 0.97 |

The histograms (figure 8) can help to validate the choice of the normalization parameter (“median” or “shorth”). Under the hypothesis that most features are not differentially expressed, each size factor represented by a red line is expected to be close to the mode of the distribution of the counts divided by their geometric means across samples.

Figure 8: Diagnostic of the estimation of the size factors.

The figure 9 shows that the scaling factors of DESeq2 and the total count normalization factors may not perform similarly.

Figure 9: Plot of the estimated size factors and the total number of reads per sample.

Boxplots are often used as a qualitative measure of the quality of the normalization process, as they show how distributions are globally affected during this process. We expect normalization to stabilize distributions across samples. Figure 10 shows boxplots of raw (left) and normalized (right) data respectively.

Figure 10: Boxplots of raw (left) and normalized (right) read counts.

### 5 Differential analysis

#### 5.1 Modelisation

DESeq2 aims at fitting one linear model per feature. For this project, the design used is counts ~ rep + group and the goal is to estimate the models’ coefficients which can be interpreted as \(\log\_2(\texttt{FC})\). These coefficients will then be tested to get p-values and adjusted p-values.

#### 5.2 Outlier detection

Model outliers are features for which at least one sample seems unrelated to the experimental or study design. For every feature and for every sample, the Cook’s distance [6] reflects how the sample matches the model. A large value of the Cook’s distance indicates an outlier count and p-values are not computed for the corresponding feature.

#### 5.3 Dispersions estimation

The DESeq2 model assumes that the count data follow a negative binomial distribution which is a robust alternative to the Poisson law when data are over-dispersed (the variance is higher than the mean). The first step of the statistical procedure is to estimate the dispersion of the data. Its purpose is to determine the shape of the mean-variance relationship. The default is to apply a GLM (Generalized Linear Model) based method (fitType=“parametric”), which can handle complex designs but may not converge in some cases. The alternative is to use fitType=“local” as described in the original paper [3] or fitType=“mean.” The parameter used for this project is fitType=“local.” Then, DESeq2 imposes a Cox Reid-adjusted profile likelihood maximization [7 and McCarthy, 2012] and uses the maximum *a posteriori* (MAP) of the dispersion [Wu, 2013].

Figure 11: Dispersion estimates (left) and diagnostic of log-normality (right).

The left panel on figure 11 shows the result of the dispersion estimation step. The x- and y-axes represent the mean count value and the estimated dispersion respectively. Black dots represent empirical dispersion estimates for each feature (from the observed counts). The red dots show the mean-variance relationship function (fitted dispersion value) as estimated by the model. The blue dots are the final estimates from the maximum *a posteriori* and are used to perform the statistical test. Blue circles (if any) point out dispersion outliers. These are features with a very high empirical variance (computed from observed counts). These high dispersion values fall far from the model estimation. For these features, the statistical test is based on the empirical variance in order to be more conservative than with the MAP dispersion. These features will have low chance to be declared significant. The figure on the right panel allows to check the hypothesis of log-normality of the dispersions.

#### 5.4 Statistical test for differential expression

Once the dispersion estimation and the model fitting have been done, DESeq2 can perform the statistical testing. Figure 12 shows the distributions of raw p-values computed by the statistical test for the comparison(s) done. This distribution is expected to be a mixture of a uniform distribution on \([0,1]\) and a peak around 0 corresponding to the differentially expressed features. If raw p-values doesn’t follow the expected distribution, FDR can be re-estimated using fdrtool [8].

#### 5.5 Independent filtering

DESeq2 can perform an independent filtering to increase the detection power of differentially expressed features at the same experiment-wide type I error. Since features with very low counts are not likely to see significant differences typically due to high dispersion, it defines a threshold on the mean of the normalized counts irrespective of the biological condition. This procedure is independent because the information about the variables in the design formula is not used [4].

Table 6 reports the thresholds used for each comparison and the number of features discarded by the independent filtering. Adjusted p-values of discarded features are then set to NA.

Table 6: Number of features discarded by the independent filtering for each comparison.

| Test vs Ref | BaseMean Threshold | # discarded |
| --- | --- | --- |
| iEF EF vs iEF empty | 2.68 | 10414 |
| iEF EE4 vs iEF empty | 3.06 | 11108 |
| iEF EE4 vs iEF EF | 9.23 | 16662 |

#### 5.6 Final results

A p-value adjustment is performed to take into account multiple testing and control the false positive rate to a chosen level \(\alpha\). For this analysis, a BH p-value adjustment was performed [9 and BY2001] and the level of False Discovery Rate (FDR) was set to 0.05.

Table 7: Number of up-, down- and total number of differentially expressed features for each comparison.

| Test vs Ref | min baseMean | # down | # up | # total |
| --- | --- | --- | --- | --- |
| iEF EF vs iEF empty | 2.69 | 4344 | 5062 | 9406 |
| iEF EE4 vs iEF empty | 3.06 | 3469 | 3810 | 7279 |
| iEF EE4 vs iEF EF | 9.24 | 2085 | 1576 | 3661 |

Table 8: View of differentially expressed features. Each table below shows top and bottom two (up- and down-regulated) significant features in each comparison. These tables can serve as a diagnostic tool to make sure significant features are indeed different in the samples compared.

Table 8.1: **iEF\_EF vs iEF\_empty(ref)**

| Id | geneSymbol | iEF\_emptyrep1 | iEF\_emptyrep2 | iEF\_EFrep1 | iEF\_EFrep2 | norm.iEF\_emptyrep1 | norm.iEF\_emptyrep2 | norm.iEF\_EFrep1 | norm.iEF\_EFrep2 | mean.norm.iEF\_empty | mean.norm.iEF\_EF | FoldChange | log2FoldChange | pvalue | padj | dispGeneEst | dispFit | dispMAP | dispersion | betaConv | maxCooks |
| --- | --- | --- | --- | --- | --- | --- | --- | --- | --- | --- | --- | --- | --- | --- | --- | --- | --- | --- | --- | --- | --- |
| ENSG00000183019 | MCEMP1 | 0 | 0 | 26 | 75 | 0 | 0 | 24 | 66 | 0 | 45 | 197.975 | 7.629 | 6.35e-07 | 4.13e-06 | 0.0640 | 0.1245 | 0.1152 | 0.1152 | TRUE | NA |
| ENSG00000117154 | IGSF21 | 1 | 3 | 275 | 523 | 1 | 3 | 251 | 462 | 2 | 356 | 162.148 | 7.341 | 6.52e-21 | 1.62e-19 | 0.1158 | 0.0300 | 0.0393 | 0.0393 | TRUE | NA |
| ENSG00000168214 | RBPJ | 3957 | 3905 | 5072 | 5795 | 4087 | 4539 | 4633 | 5117 | 4313 | 4875 | 1.130 | 0.177 | 1.29e-02 | 3.67e-02 | 0.0000 | 0.0039 | 0.0022 | 0.0022 | TRUE | NA |
| ENSG00000101639 | CEP192 | 4475 | 4022 | 5738 | 5942 | 4622 | 4674 | 5241 | 5246 | 4648 | 5244 | 1.128 | 0.174 | 1.33e-02 | 3.77e-02 | 0.0000 | 0.0038 | 0.0022 | 0.0022 | TRUE | NA |
| ENSG00000082482 | KCNK2 | 461 | 1604 | 0 | 1 | 476 | 1864 | 0 | 1 | 1170 | 0 | -2340.154 | -11.192 | 1.08e-19 | 2.49e-18 | 0.0956 | 0.0134 | 0.0166 | 0.0166 | TRUE | NA |
| ENSG00000111341 | MGP | 10995 | 22608 | 8 | 39 | 11357 | 26276 | 7 | 34 | 18816 | 20 | -1003.036 | -9.970 | 1.85e-136 | 8.71e-134 | 0.0505 | 0.0037 | 0.0125 | 0.0505 | TRUE | NA |
| ENSG00000168137 | SETD5 | 7377 | 7765 | 7291 | 9156 | 7620 | 9025 | 6660 | 8084 | 8322 | 7372 | -1.130 | -0.176 | 1.46e-02 | 4.07e-02 | 0.0006 | 0.0037 | 0.0024 | 0.0024 | TRUE | NA |
| ENSG00000164828 | SUN1 | 6270 | 6544 | 6498 | 7447 | 6476 | 7606 | 5935 | 6575 | 7041 | 6255 | -1.123 | -0.168 | 1.85e-02 | 4.99e-02 | 0.0004 | 0.0037 | 0.0023 | 0.0023 | TRUE | NA |

Table 8.2: **iEF\_EE4 vs iEF\_empty(ref)**

| Id | geneSymbol | iEF\_emptyrep1 | iEF\_emptyrep2 | iEF\_EE4rep1 | iEF\_EE4rep2 | norm.iEF\_emptyrep1 | norm.iEF\_emptyrep2 | norm.iEF\_EE4rep1 | norm.iEF\_EE4rep2 | mean.norm.iEF\_empty | mean.norm.iEF\_EE4 | FoldChange | log2FoldChange | pvalue | padj | dispGeneEst | dispFit | dispMAP | dispersion | betaConv | maxCooks |
| --- | --- | --- | --- | --- | --- | --- | --- | --- | --- | --- | --- | --- | --- | --- | --- | --- | --- | --- | --- | --- | --- |
| ENSG00000124233 | SEMG1 | 1 | 1 | 221 | 131 | 1 | 1 | 210 | 135 | 1 | 172 | 163.865 | 7.356 | 5.92e-12 | 1.08e-10 | 0.0000 | 0.0440 | 0.0446 | 0.0446 | TRUE | NA |
| ENSG00000225899 | FRG2B | 0 | 2 | 168 | 193 | 0 | 2 | 160 | 200 | 1 | 180 | 159.985 | 7.322 | 1.60e-03 | 7.29e-03 | 2.0951 | 0.0480 | 0.0537 | 2.0951 | TRUE | NA |
| ENSG00000116237 | ICMT | 5882 | 5105 | 6965 | 6739 | 6076 | 5933 | 6617 | 6967 | 6004 | 6792 | 1.131 | 0.177 | 1.47e-02 | 4.98e-02 | 0.0005 | 0.0037 | 0.0024 | 0.0024 | TRUE | NA |
| ENSG00000198763 | MT-ND2 | 16176 | 10084 | 19856 | 12709 | 16709 | 11720 | 18864 | 13140 | 14214 | 16002 | 1.125 | 0.170 | 1.39e-02 | 4.74e-02 | 0.0003 | 0.0038 | 0.0022 | 0.0022 | TRUE | NA |
| ENSG00000082482 | KCNK2 | 461 | 1604 | 0 | 1 | 476 | 1864 | 0 | 1 | 1170 | 0 | -2085.999 | -11.027 | 1.06e-18 | 3.37e-17 | 0.0956 | 0.0134 | 0.0166 | 0.0166 | TRUE | NA |
| ENSG00000137441 | FGFBP2 | 709 | 3416 | 1 | 3 | 732 | 3970 | 1 | 3 | 2351 | 2 | -1149.836 | -10.167 | 3.77e-43 | 3.33e-41 | 0.0096 | 0.0084 | 0.0084 | 0.0084 | TRUE | NA |
| ENSG00000048828 | FAM120A | 6968 | 5394 | 6630 | 5391 | 7197 | 6269 | 6299 | 5574 | 6733 | 5936 | -1.134 | -0.181 | 1.13e-02 | 3.99e-02 | 0.0000 | 0.0037 | 0.0023 | 0.0023 | TRUE | NA |
| ENSG00000082641 | NFE2L1 | 9203 | 8810 | 8652 | 8974 | 9506 | 10239 | 8220 | 9278 | 9872 | 8749 | -1.130 | -0.176 | 1.15e-02 | 4.06e-02 | 0.0000 | 0.0037 | 0.0022 | 0.0022 | TRUE | NA |

Table 8.3: **iEF\_EE4 vs iEF\_EF(ref)**

| Id | geneSymbol | iEF\_EFrep1 | iEF\_EFrep2 | iEF\_EE4rep1 | iEF\_EE4rep2 | norm.iEF\_EFrep1 | norm.iEF\_EFrep2 | norm.iEF\_EE4rep1 | norm.iEF\_EE4rep2 | mean.norm.iEF\_EF | mean.norm.iEF\_EE4 | FoldChange | log2FoldChange | pvalue | padj | dispGeneEst | dispFit | dispMAP | dispersion | betaConv | maxCooks |
| --- | --- | --- | --- | --- | --- | --- | --- | --- | --- | --- | --- | --- | --- | --- | --- | --- | --- | --- | --- | --- | --- |
| ENSG00000100678 | SLC8A3 | 0 | 0 | 43 | 85 | 0 | 0 | 41 | 88 | 0 | 64 | 362.522 | 8.502 | 2.03e-08 | 5.62e-07 | 0.0000 | 0.1015 | 0.0950 | 0.0950 | TRUE | NA |
| ENSG00000225899 | FRG2B | 3 | 0 | 168 | 193 | 3 | 0 | 160 | 200 | 2 | 180 | 128.516 | 7.006 | 1.80e-03 | 1.32e-02 | 2.0951 | 0.0480 | 0.0537 | 2.0951 | TRUE | NA |
| ENSG00000153250 | RBMS1 | 6042 | 5656 | 6540 | 5570 | 5519 | 4994 | 6213 | 5759 | 5256 | 5986 | 1.139 | 0.188 | 8.43e-03 | 4.54e-02 | 0.0000 | 0.0037 | 0.0023 | 0.0023 | TRUE | NA |
| ENSG00000139218 | SCAF11 | 6527 | 7445 | 7161 | 7190 | 5962 | 6574 | 6803 | 7434 | 6268 | 7118 | 1.136 | 0.184 | 7.81e-03 | 4.26e-02 | 0.0000 | 0.0037 | 0.0022 | 0.0022 | TRUE | NA |
| ENSG00000117154 | IGSF21 | 275 | 523 | 0 | 1 | 251 | 462 | 0 | 1 | 356 | 0 | -702.665 | -9.457 | 5.74e-12 | 2.62e-10 | 0.1158 | 0.0300 | 0.0393 | 0.0393 | TRUE | NA |
| ENSG00000171201 | SMR3B | 7 | 21 | 0 | 0 | 6 | 19 | 0 | 0 | 12 | 0 | -62.318 | -5.962 | 1.30e-04 | 1.43e-03 | 0.0360 | 0.1385 | 0.1247 | 0.1247 | TRUE | NA |
| ENSG00000083312 | TNPO1 | 18361 | 16659 | 15173 | 12830 | 16771 | 14709 | 14415 | 13265 | 15740 | 13840 | -1.136 | -0.184 | 9.23e-03 | 4.87e-02 | 0.0007 | 0.0038 | 0.0023 | 0.0023 | TRUE | NA |
| ENSG00000056097 | ZFR | 10116 | 10351 | 8632 | 7787 | 9240 | 9139 | 8201 | 8051 | 9190 | 8126 | -1.131 | -0.178 | 7.43e-03 | 4.10e-02 | 0.0000 | 0.0037 | 0.0020 | 0.0020 | TRUE | NA |

Figure 13 represents the MA-plot of the data for the comparisons done, where differentially expressed features are highlighted in red. A MA-plot represents the log ratio of differential expression as a function of the mean intensity for each feature. Triangles correspond to features having a too low/high \(\log\_2(\text{FC})\) to be displayed on the plot.

Figure 13: MA-plot(s) of each comparison. Red dots represent significantly differentially expressed features.

Figure 14 shows the volcano plots for the comparisons performed and differentially expressed features are still highlighted in red. A volcano plot represents the log of the adjusted P value as a function of the log ratio of differential expression.

Figure 14: Volcano plot(s) of each comparison. Red dots represent significantly differentially expressed features.

Heatmaps showing the rlog normalized counts of up-/down-regulated genes in each sample can be found in figures/diffPlot.pdf. This plot again can serve as a diagnostic tool to make sure the normalized counts in all of the replicates are higher/lower compared to control for up-/down-regulated genes, respectively.

Full results as well as lists of differentially expressed features are provided in the following text files which can be easily read in a spreadsheet. For each comparison:

- TestVsRef.complete.txt contains results for all the features;
- TestVsRef.up.txt contains results for significantly up-regulated features. Features are ordered from the most significant adjusted p-value to the less significant one;
- TestVsRef.down.txt contains results for significantly down-regulated features. Features are ordered from the most significant adjusted p-value to the less significant one.

These files contain the following columns:

- Id: unique feature identifier;
- sampleName: raw counts per sample;
- norm.sampleName: rounded normalized counts per sample;
- baseMean: base mean over all samples;
- iEF\_empty, iEF\_EF and iEF\_EE4: means (rounded) of normalized counts of the biological conditions;
- FoldChange: fold change of expression, calculated as \(2^{\log\_2(\text{FC})}\);
- log2FoldChange: \(\log\_2(\text{FC})\) as estimated by the GLM model. It reflects the differential expression between Test and Ref and can be interpreted as \(\log\_2(\frac{\text{Test}}{\text{Ref}})\). If this value is:
  - around 0: the feature expression is similar in both conditions;
  - positive: the feature is up-regulated (\(\text{Test} > \text{Ref}\));
  - negative: the feature is down-regulated (\(\text{Test} < \text{Ref}\));
- stat: Wald statistic for the coefficient tested;
- pvalue: raw p-value from the statistical test;
- padj: adjusted p-value on which the cut-off \(\alpha\) is applied;
- dispGeneEst: dispersion parameter estimated from feature counts (i.e. black dots on figure 11);
- dispFit: dispersion parameter estimated from the model (i.e. red dots on figure 11);
- dispMAP: dispersion parameter estimated from the Maximum *A Posteriori* model;
- dispersion: final dispersion parameter used to perform the test (i.e. blue dots and circles on figure 11);
- betaConv: convergence of the coefficients of the model (TRUE or FALSE);
- maxCooks: maximum Cook’s distance of the feature.

### 6 R session information and parameters

The versions of the R software and Bioconductor packages used for this analysis are listed below. It is important to save them if one wants to re-perform the analysis in the same conditions.

- R version 4.0.3 (2020-10-10), x86\_64-conda-linux-gnu
- Locale: LC\_CTYPE=en\_US.UTF-8, LC\_NUMERIC=C, LC\_TIME=en\_US.UTF-8, LC\_COLLATE=en\_US.UTF-8, LC\_MONETARY=en\_US.UTF-8, LC\_MESSAGES=en\_US.UTF-8, LC\_PAPER=en\_US.UTF-8, LC\_NAME=C, LC\_ADDRESS=C, LC\_TELEPHONE=C, LC\_MEASUREMENT=en\_US.UTF-8, LC\_IDENTIFICATION=C
- Running under: CentOS Linux 7 (Core)
- Matrix products: default
- BLAS/LAPACK: /opt/envs/sartools\_env/lib/libopenblasp-r0.3.10.so
- Base packages: base, datasets, graphics, grDevices, grid, methods, parallel, stats, stats4, utils
- Other packages: AnnotationDbi 1.52.0, AnnotationFilter 1.14.0, Biobase 2.50.0, BiocGenerics 0.36.0, DESeq2 1.30.0, edgeR 3.32.0, EnsDb.Hsapiens.v86 2.99.0, ensembldb 2.14.0, fdrtool 1.2.15, GenomeInfoDb 1.26.1, GenomicFeatures 1.42.1, GenomicRanges 1.42.0, ggplot2 3.3.2, gridExtra 2.3, IRanges 2.24.0, kableExtra 1.3.1, limma 3.46.0, MatrixGenerics 1.2.0, matrixStats 0.57.0, pheatmap 1.0.12, Polychrome 1.2.6, RColorBrewer 1.1-2, S4Vectors 0.28.0, SARTools 1.7.3, SummarizedExperiment 1.20.0, tibble 3.0.4
- Loaded via a namespace (and not attached): annotate 1.68.0, askpass 1.1, assertthat 0.2.1, BiocFileCache 1.14.0, BiocParallel 1.24.1, biomaRt 2.46.0, Biostrings 2.58.0, bit 4.0.4, bit64 4.0.5, bitops 1.0-6, blob 1.2.1, cli 2.2.0, colorspace 2.0-0, compiler 4.0.3, crayon 1.3.4, curl 4.3, DBI 1.1.0, dbplyr 2.0.0, DelayedArray 0.16.0, digest 0.6.27, dplyr 1.0.2, ellipsis 0.3.1, evaluate 0.14, fansi 0.4.1, farver 2.0.3, fastmap 1.1.0, genefilter 1.72.0, geneplotter 1.68.0, generics 0.1.0, GenomeInfoDbData 1.2.4, GenomicAlignments 1.26.0, GGally 2.0.0, ggdendro 0.1.22, ggrepel 0.8.2, glue 1.4.2, gtable 0.3.0, highr 0.8, hms 0.5.3, htmltools 0.5.7, httr 1.4.2, knitr 1.30, labeling 0.4.2, lattice 0.20-41, lazyeval 0.2.2, lifecycle 0.2.0, locfit 1.5-9.4, magrittr 2.0.1, MASS 7.3-53, Matrix 1.2-18, memoise 1.1.0, munsell 0.5.0, openssl 1.4.3, pillar 1.4.7, pkgconfig 2.0.3, plyr 1.8.6, prettyunits 1.1.1, progress 1.2.2, ProtGenerics 1.22.0, purrr 0.3.4, R6 2.5.0, rappdirs 0.3.1, Rcpp 1.0.5, RCurl 1.98-1.2, reshape 0.8.8, rlang 1.0.6, rmarkdown 2.5, Rsamtools 2.6.0, RSQLite 2.2.1, rstudioapi 0.13, rtracklayer 1.50.0, rvest 0.3.6, scales 1.1.1, scatterplot3d 0.3-41, splines 4.0.3, stringi 1.5.3, stringr 1.4.0, survival 3.2-7, tidyselect 1.1.0, tools 4.0.3, vctrs 0.3.6, viridisLite 0.4.0, webshot 0.5.2, withr 2.3.0, xfun 0.19, XML 3.99-0.5, xml2 1.3.2, xtable 1.8-4, XVector 0.30.0, yaml 2.2.1, zlibbioc 1.36.0

Parameter values used for this analysis are:

- workDir: /mnt/outputs/diff\_analysis\_rslt
- projectName: RNA-seq\_differential\_analysis
- author: NA
- targetFile: target.txt
- rawDir: ../STAR\_2pass/Pass2
- featuresToRemove: alignment\_not\_unique, ambiguous, no\_feature, not\_aligned, too\_low\_aQual
- varInt: group
- condRef: iEF\_empty, iEF\_empty, iEF\_empty, iEF\_empty
- batch: rep
- fitType: local
- cooksCutoff: TRUE
- independentFiltering: TRUE
- alpha: 0.05
- pAdjustMethod: BH
- typeTrans: VST
- locfunc: median
- colors: #ADD8E6, #FFA500, #C71585

### Bibliography

1.

R Core Team. *R: A language and environment for statistical computing*. Vienna, Austria : R Foundation for Statistical Computing, 2017 :

2.

Gentleman RC, Carey VJ, Bates DM, *et al.* Bioconductor: Open software development for computational biology and bioinformatics. *Genome Biology* 2004 ; 5 : R80.

3.

Anders S, Huber W. Differential expression analysis for sequence count data. *Genome Biology* 2010 ; 11 : R106.

4.

Love MI, Huber W, Anders S. Moderated estimation of fold change and dispersion for RNA-seq data with DESeq2. *Genome Biology* 2014 ; 15 : 550.

5.

Schulze SK, Kanwar R, Gölzenleuchter M, *et al.* SERE: Single-parameter quality control and sample comparison for RNA-seq. *BMC Genomics* 2012 ; 13 : 524.

6.

Cook RD. Detection of influential observation in linear regression. *Technometrics* 1977 ; 19 : 15–18.

7.

Cox DR, Reid N. Parameter orthogonality and approximate conditional inference. *Journal of the Royal Statistical Society. Series B (Methodological)* 1987 ; 49 : 1–39.

8.

Strimmer K. fdrtool: a versatile R package for estimating local and tail area-based false discovery rates. *Bioinformatics* 2008 ; 24 : 1461–1462.

9.

Benjamini Y, Hochberg Y. Controlling the false discovery rate: A practical and powerful approach to multiple testing. *Journal of the Royal Statistical Society. Series B (Methodological)* 1995 ; 57 : 289–300.
