## Supplementary File 3 for "RNA-SeqEZPZ: A Point-and-Click Pipeline for Comprehensive Transcriptomics Analysis with Interactive Visualizations"

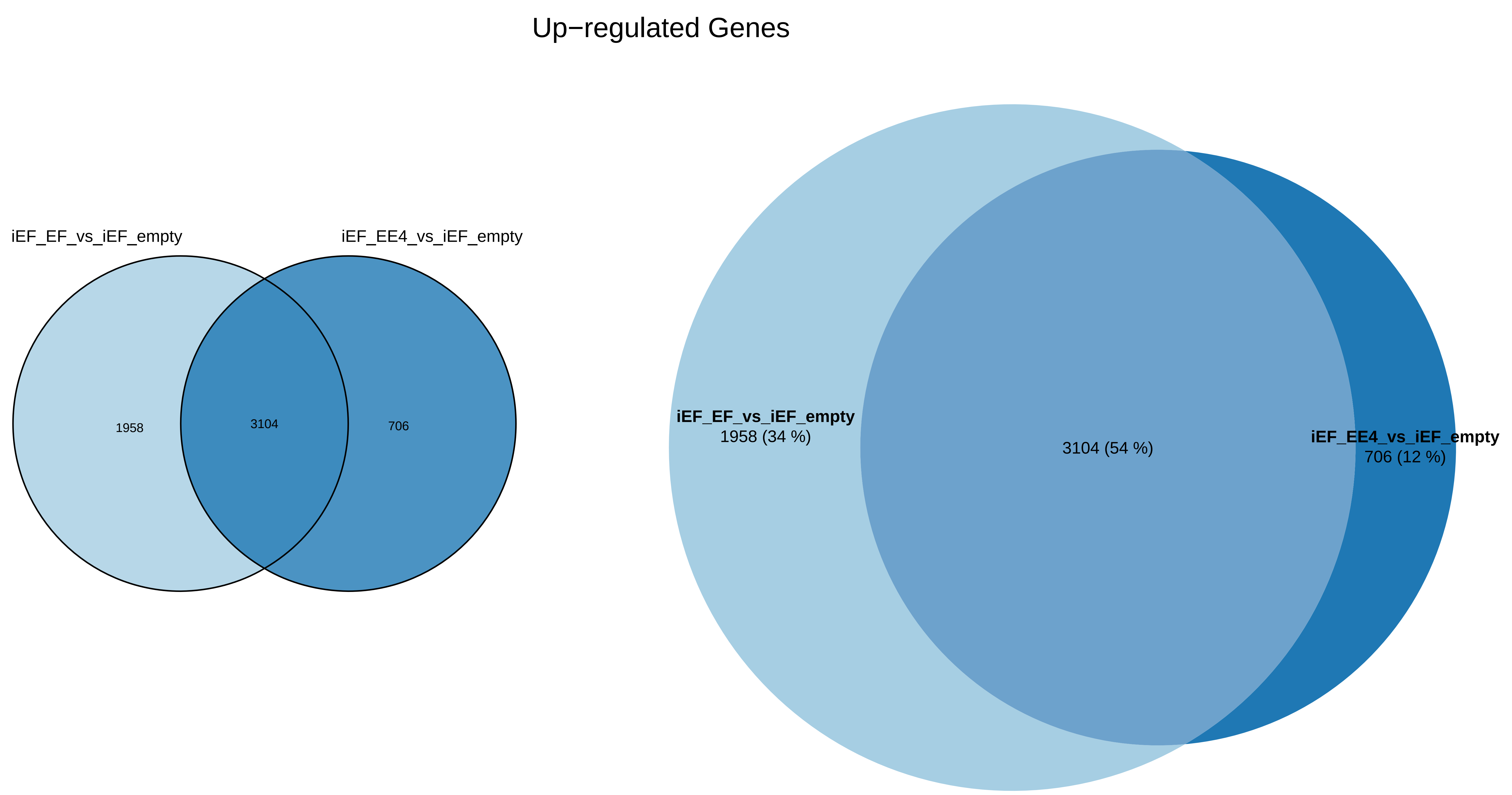
Up−regulated Genes
iEF_EF_vs_iEF_empty
iEF_EE4_vs_iEF_empty
1958
3104
706
iEF_EF_vs_iEF_empty
1958 (34 %)
3104 (54 %)
iEF_EE4_vs_iEF_empty
706 (12 %)

Down−regulated Genes

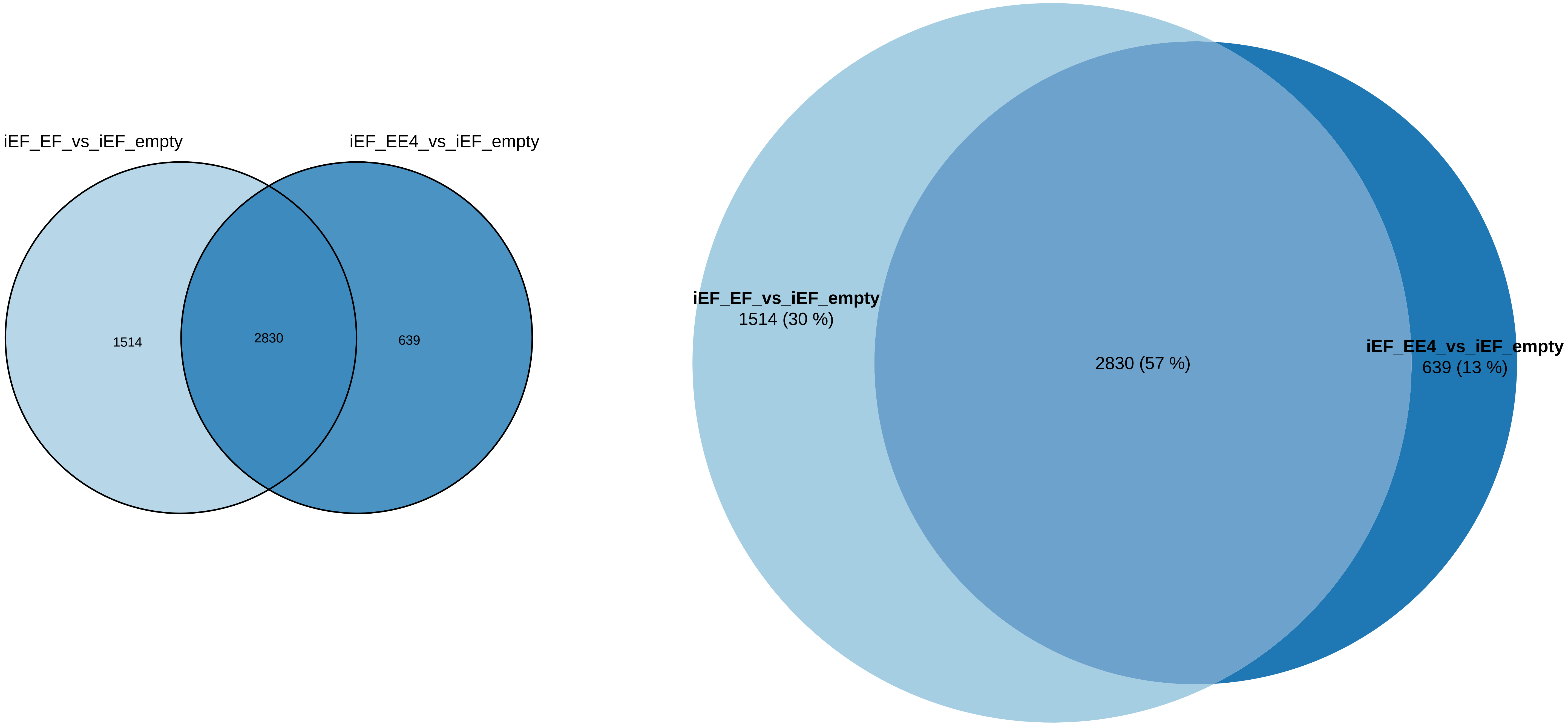

### Significance of overlaps

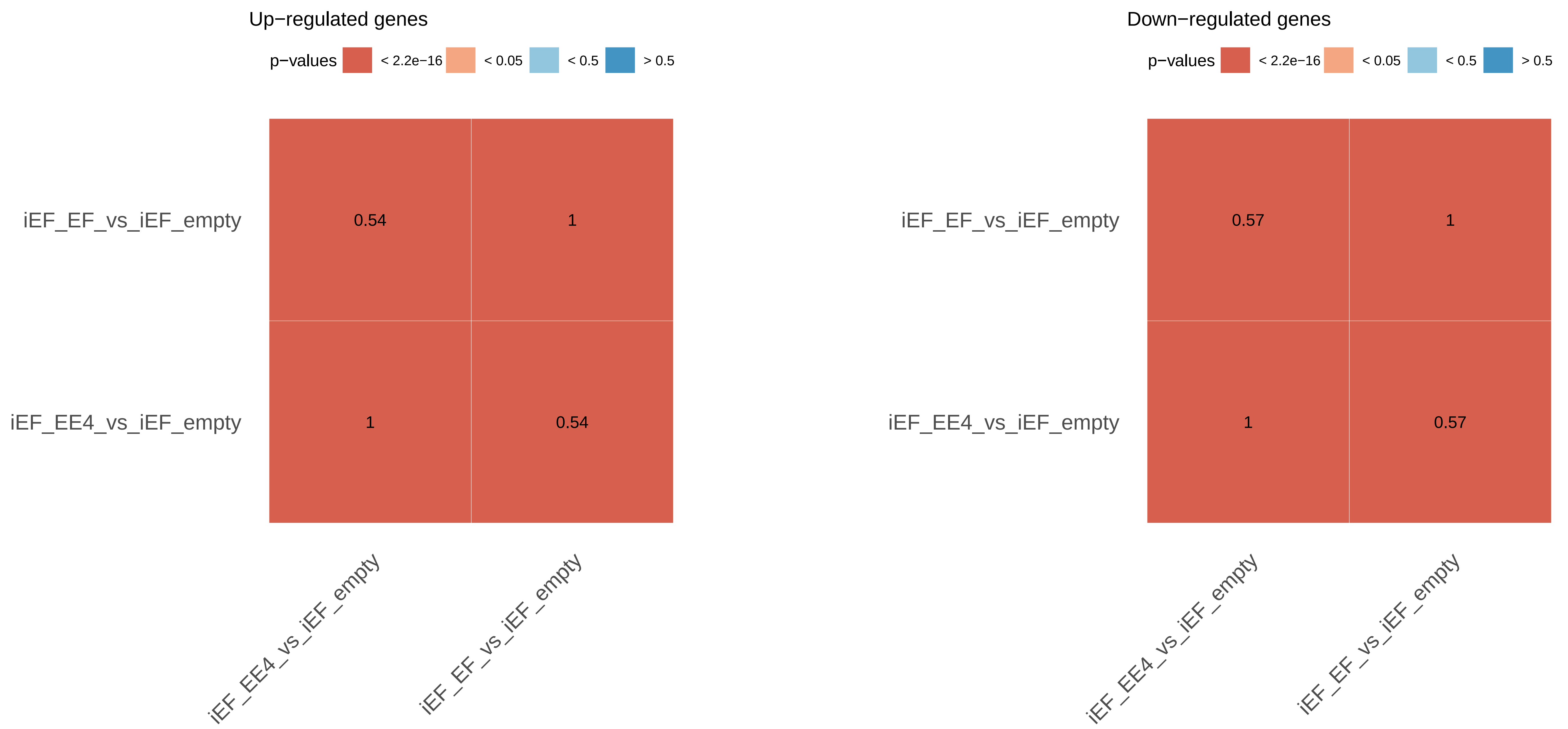


Color represents p−values of overlap.
Number inside a cell is Jaccard similarity index.
0 means no similarity, 1 means identical.

Color represents p−values of overlap.
Number inside a cell is Jaccard similarity index.
0 means no similarity, 1 means identical.

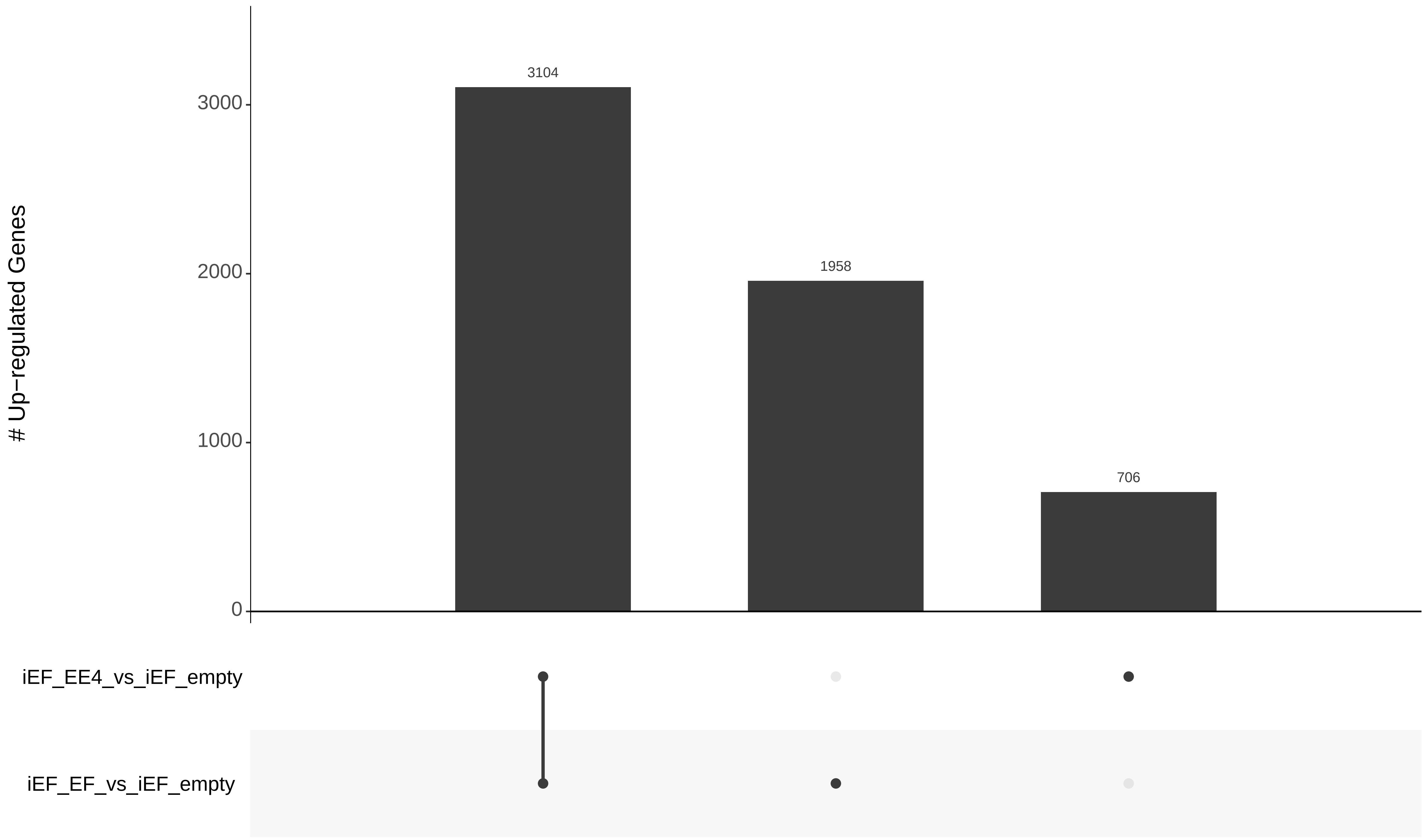

### Up−regulated Genes
3000
2000
1000
0
3104
1958
706
iEF_EE4_vs_iEF_empty
iEF_EF_vs_iEF_empty

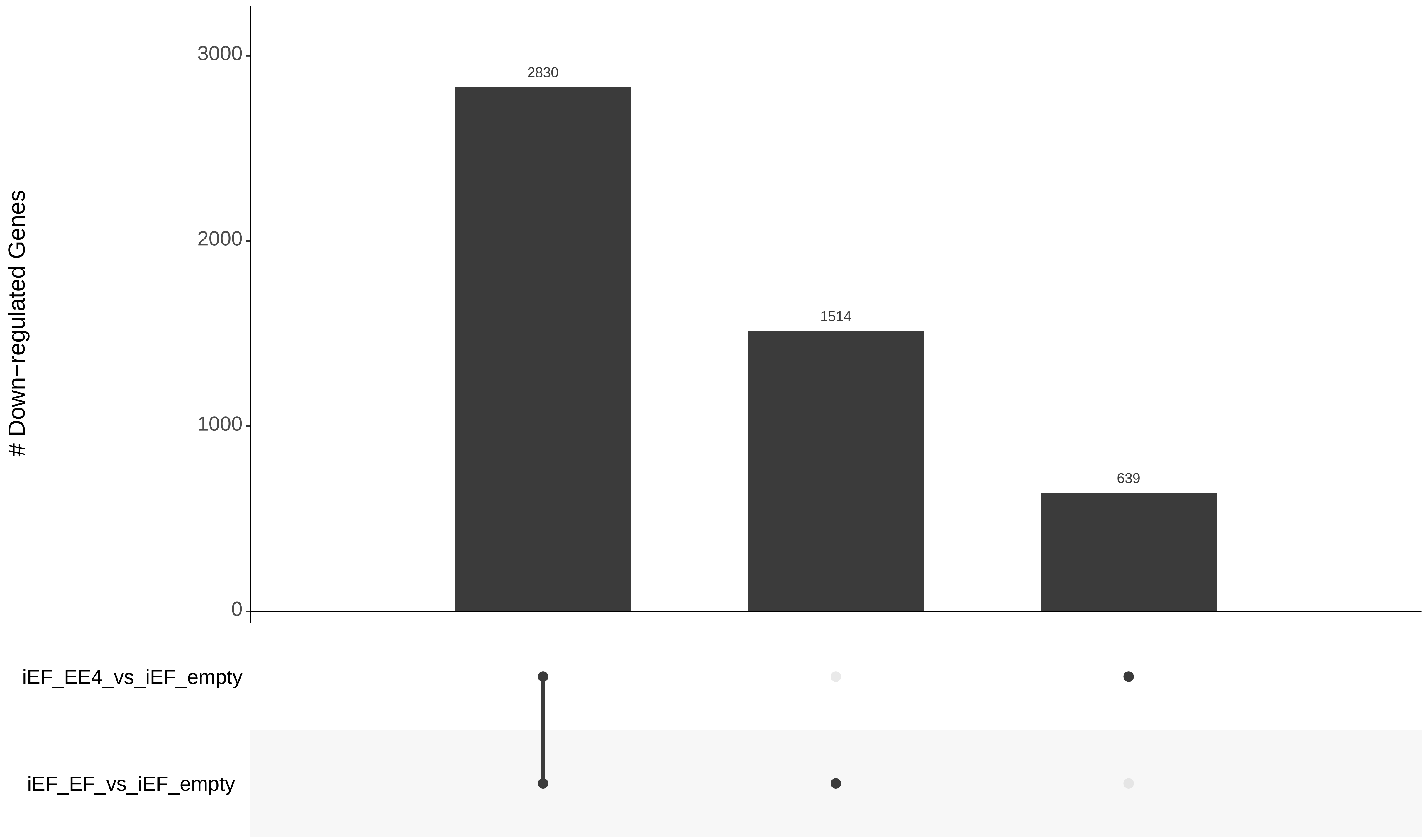

### Down−regulated Genes
3000
2000
1000
0
2830
1514
639
iEF_EE4_vs_iEF_empty
iEF_EF_vs_iEF_empty

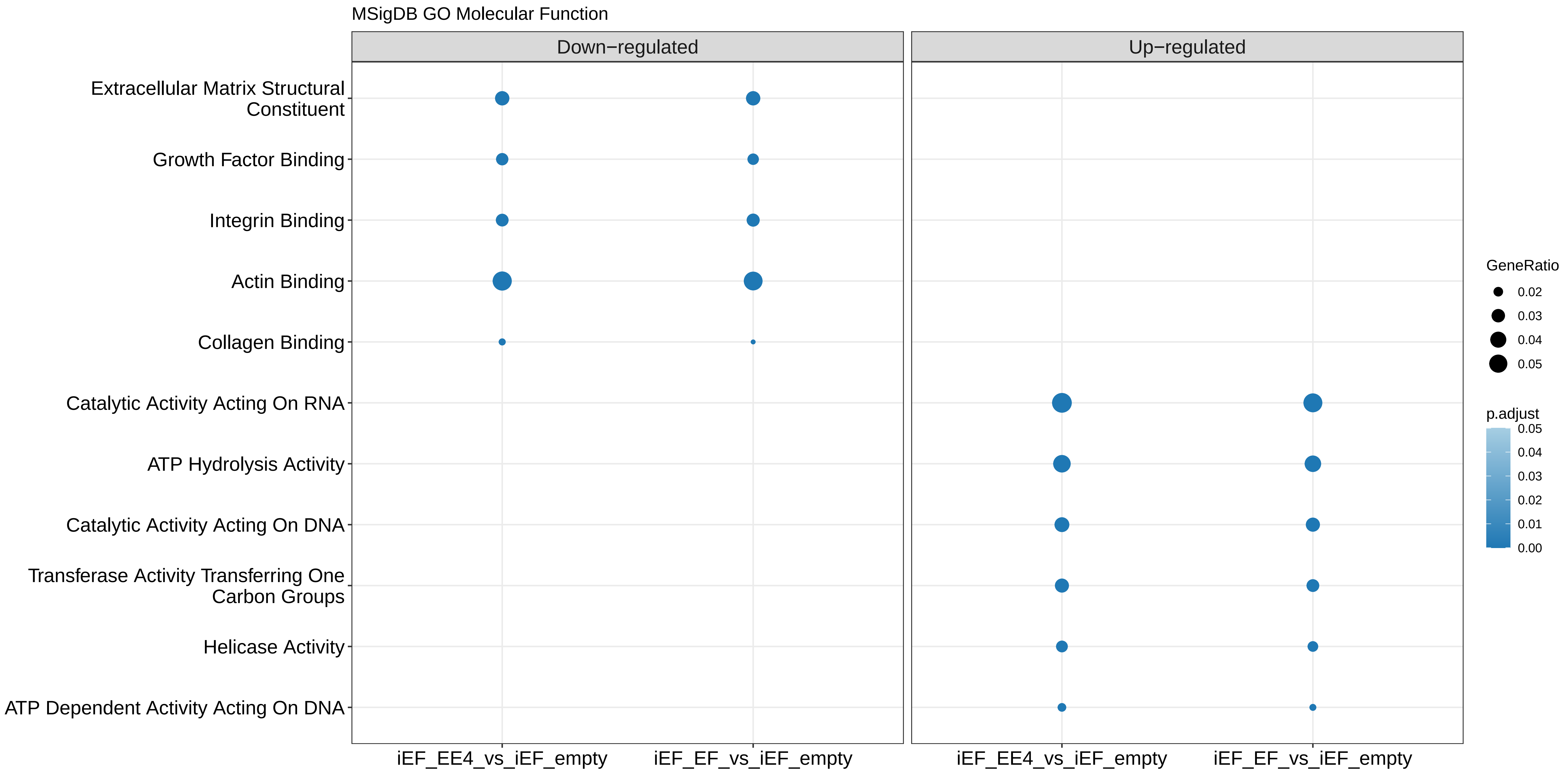

MSigDB GO Molecular Function
Down−regulated
Up−regulated
Extracellular Matrix Structural Constituent
Growth Factor Binding
Integrin Binding
Actin Binding
Collagen Binding
Catalytic Activity Acting On RNA
ATP Hydrolysis Activity
Catalytic Activity Acting On DNA
Transferase Activity Transferring One Carbon Groups
Helicase Activity
ATP Dependent Activity Acting On DNA
iEF_EE4_vs_iEF_empty
iEF_EF_vs_iEF_empty
iEF_EE4_vs_iEF_empty
iEF_EF_vs_iEF_empty
GeneRatio
0.02
0.03
0.04
0.05
p.adjust
0.05
0.04
0.03
0.02
0.01
0.00

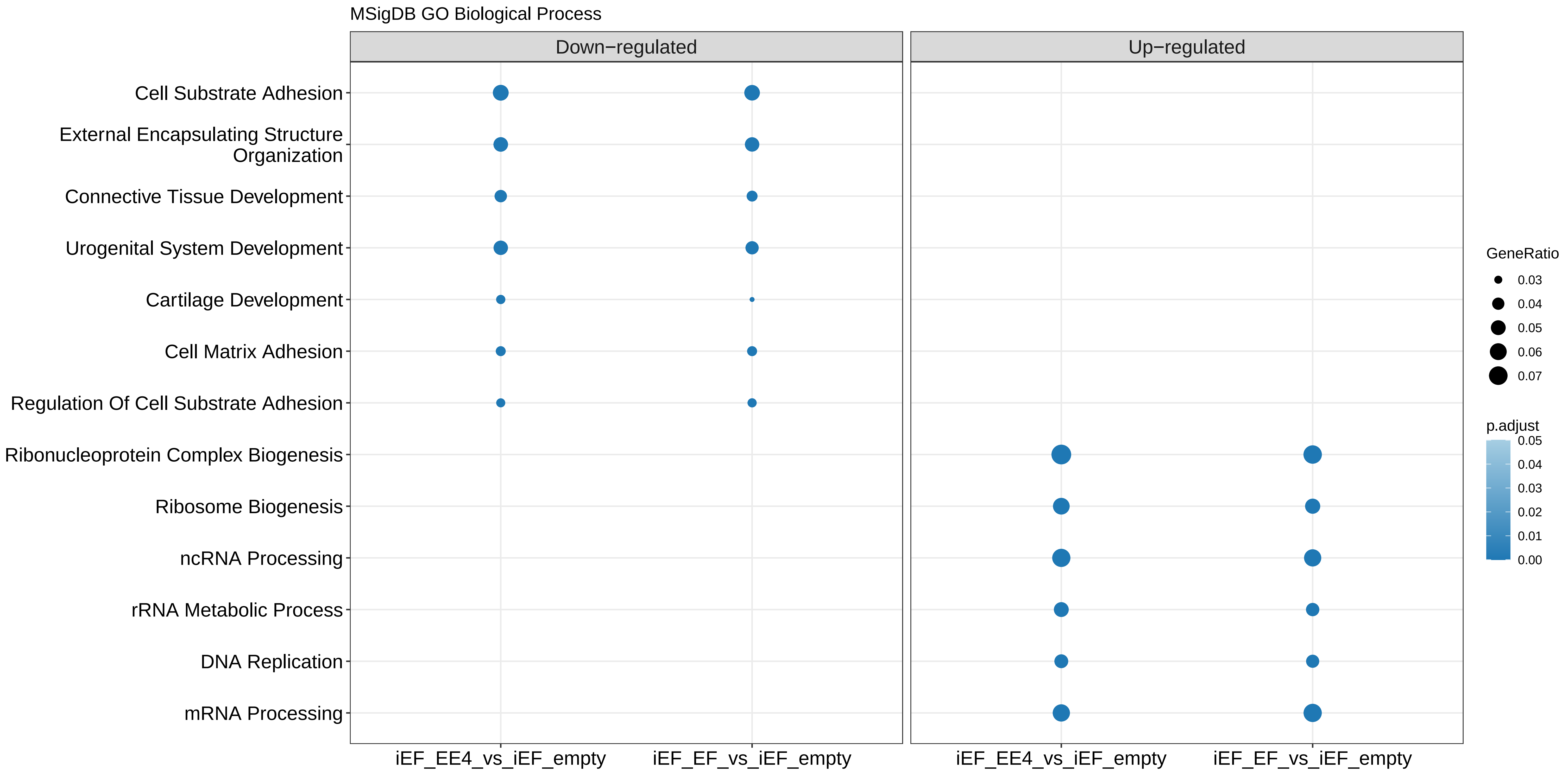
MSigDB GO Biological Process
Down−regulated
Up−regulated
Cell Substrate Adhesion
External Encapsulating Structure Organization
Connective Tissue Development
Urogenital System Development
Cartilage Development
Cell Matrix Adhesion
Regulation Of Cell Substrate Adhesion
Ribonucleoprotein Complex Biogenesis
Ribosome Biogenesis
ncRNA Processing
rRNA Metabolic Process
DNA Replication
mRNA Processing
iEF_EE4_vs_iEF_empty
iEF_EF_vs_iEF_empty
iEF_EE4_vs_iEF_empty
iEF_EF_vs_iEF_empty
GeneRatio
0.03
0.04
0.05
0.06
0.07
p.adjust
0.05
0.04
0.03
0.02
0.01
0.00

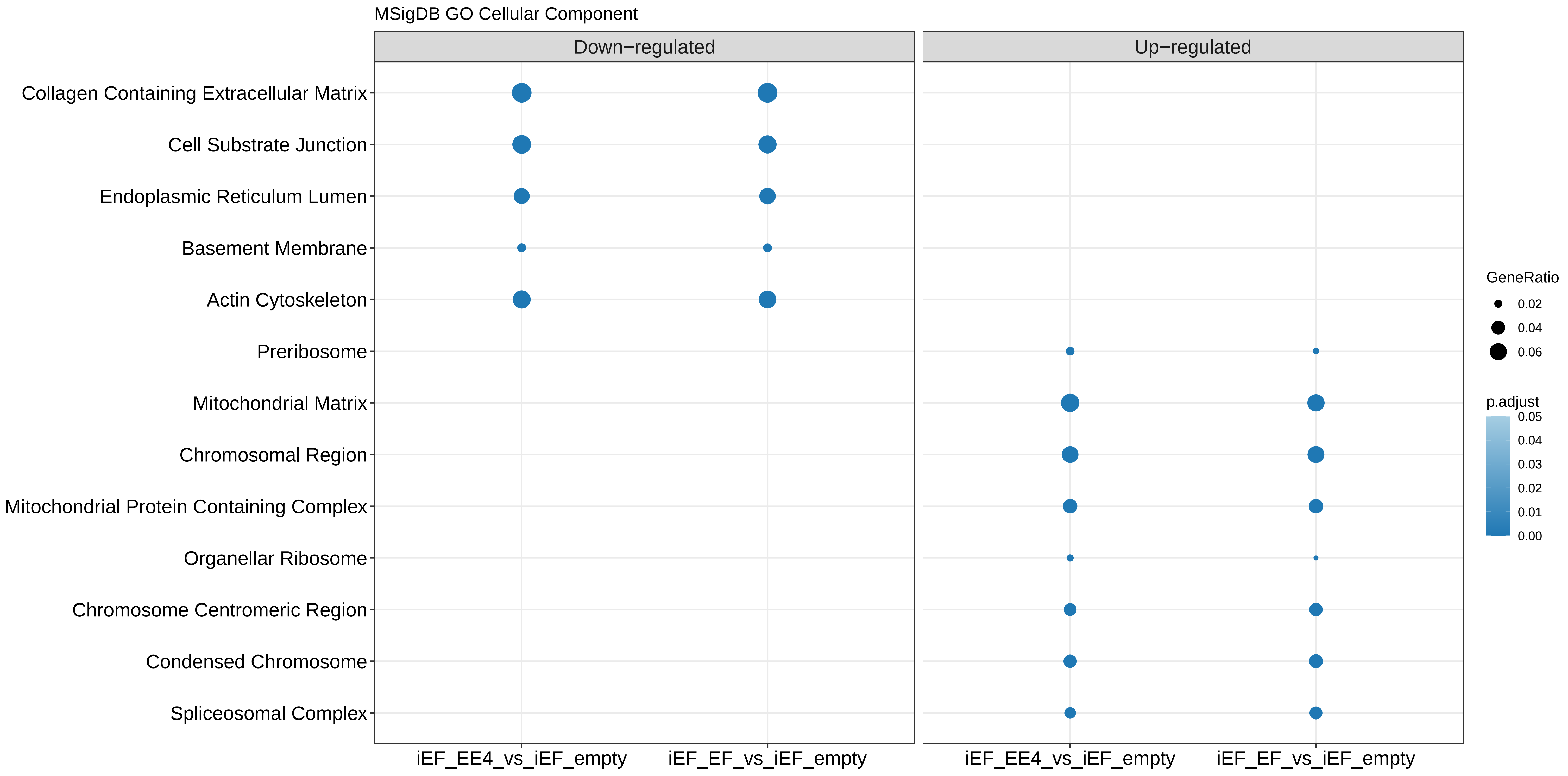

MSigDB GO Cellular Component
Down−regulated
Up−regulated
Collagen Containing Extracellular Matrix
Cell Substrate Junction
Endoplasmic Reticulum Lumen
Basement Membrane
Actin Cytoskeleton
Preribosome
Mitochondrial Matrix
Chromosomal Region
Mitochondrial Protein Containing Complex
Organellar Ribosome
Chromosome Centromeric Region
Condensed Chromosome
Spliceosomal Complex
iEF_EE4_vs_iEF_empty
iEF_EF_vs_iEF_empty
iEF_EE4_vs_iEF_empty
iEF_EF_vs_iEF_empty
GeneRatio
0.02
0.04
0.06
p.adjust
0.05
0.04
0.03
0.02
0.01
0.00

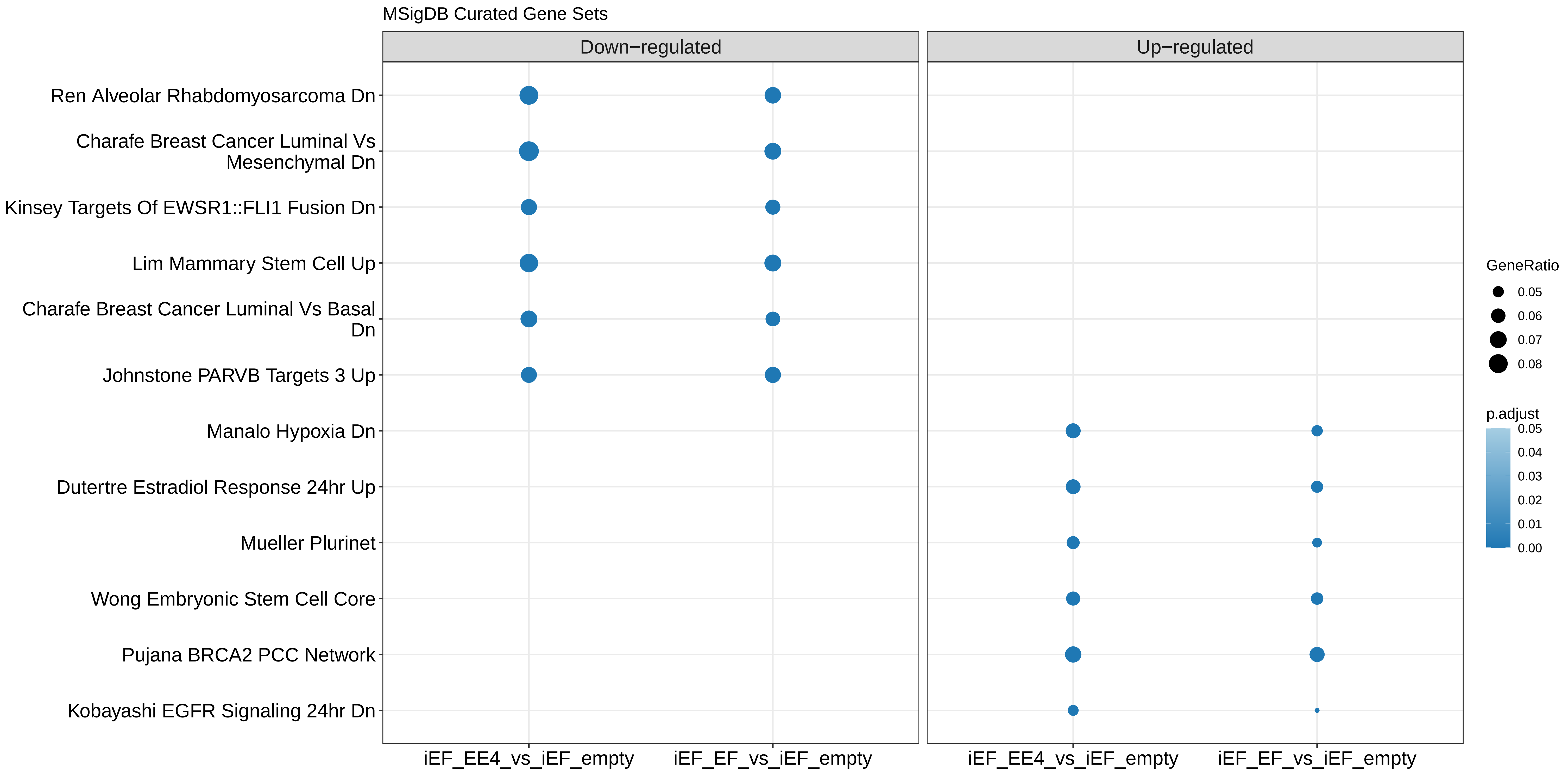

MSigDB Curated Gene Sets
Down−regulated
Up−regulated
Ren Alveolar Rhabdomyosarcoma Dn
Charafe Breast Cancer Luminal Vs Mesenchymal Dn
Kinsey Targets Of EWSR1::FLI1 Fusion Dn
Lim Mammary Stem Cell Up
Charafe Breast Cancer Luminal Vs Basal Dn
Johnstone PARVB Targets 3 Up
Manalo Hypoxia Dn
Dutertre Estradiol Response 24hr Up
Mueller Plurinet
Wong Embryonic Stem Cell Core
Pujana BRCA2 PCC Network
Kobayashi EGFR Signaling 24hr Dn
iEF_EE4_vs_iEF_empty
iEF_EF_vs_iEF_empty
iEF_EE4_vs_iEF_empty
iEF_EF_vs_iEF_empty
GeneRatio
0.05
0.06
0.07
0.08
p.adjust
0.05
0.04
0.03
0.02
0.01
0.00

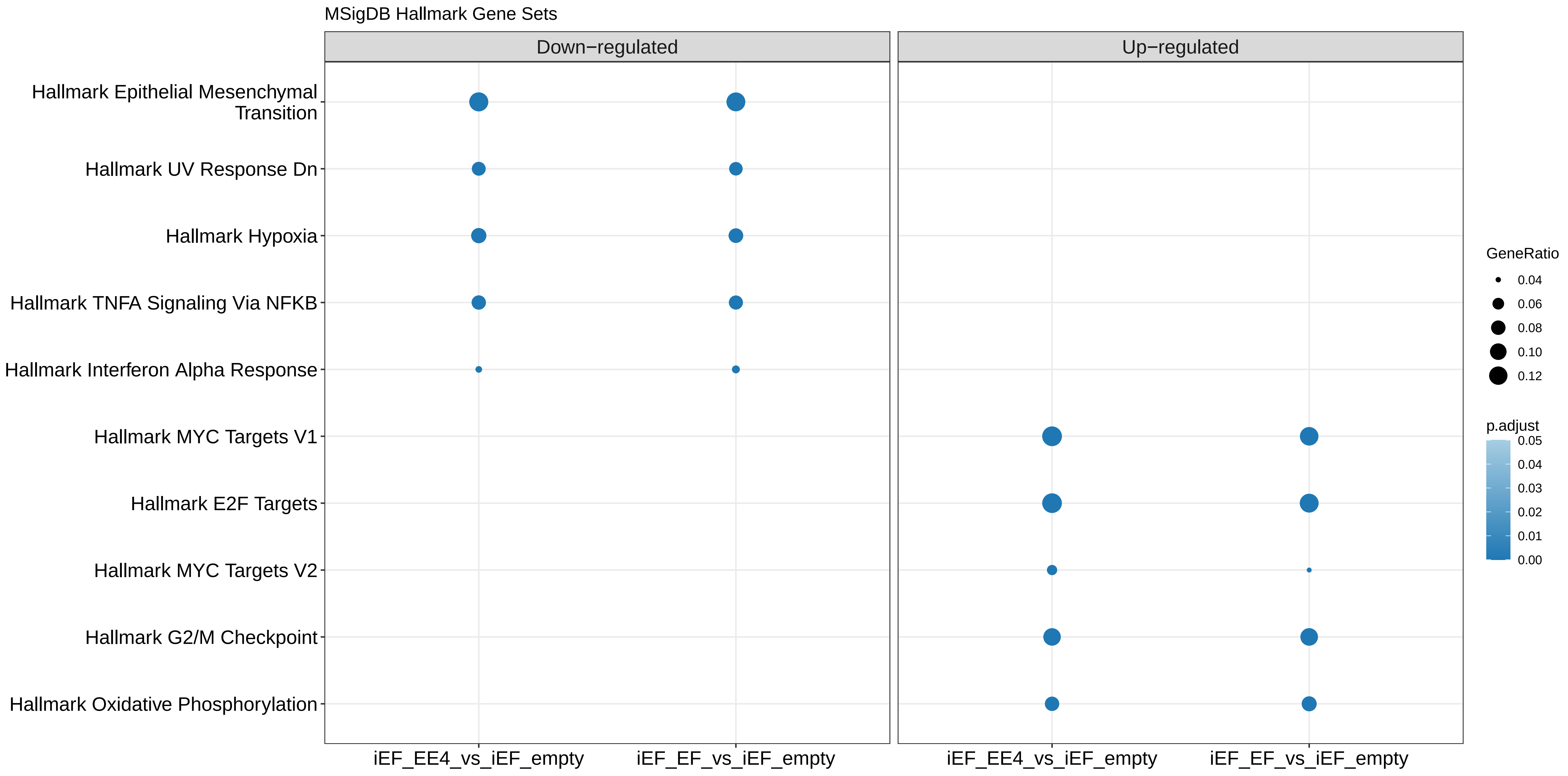

MSigDB Hallmark Gene Sets
Down−regulated
Up−regulated
Hallmark Epithelial Mesenchymal Transition
Hallmark UV Response Dn
Hallmark Hypoxia
Hallmark TNFA Signaling Via NFKB
Hallmark Interferon Alpha Response
Hallmark MYC Targets V1
Hallmark E2F Targets
Hallmark MYC Targets V2
Hallmark G2/M Checkpoint
Hallmark Oxidative Phosphorylation
iEF_EE4_vs_iEF_empty
iEF_EF_vs_iEF_empty
iEF_EE4_vs_iEF_empty
iEF_EF_vs_iEF_empty
GeneRatio
0.04
0.06
0.08
0.10
0.12
p.adjust
0.05
0.04
0.03
0.02
0.01
0.00

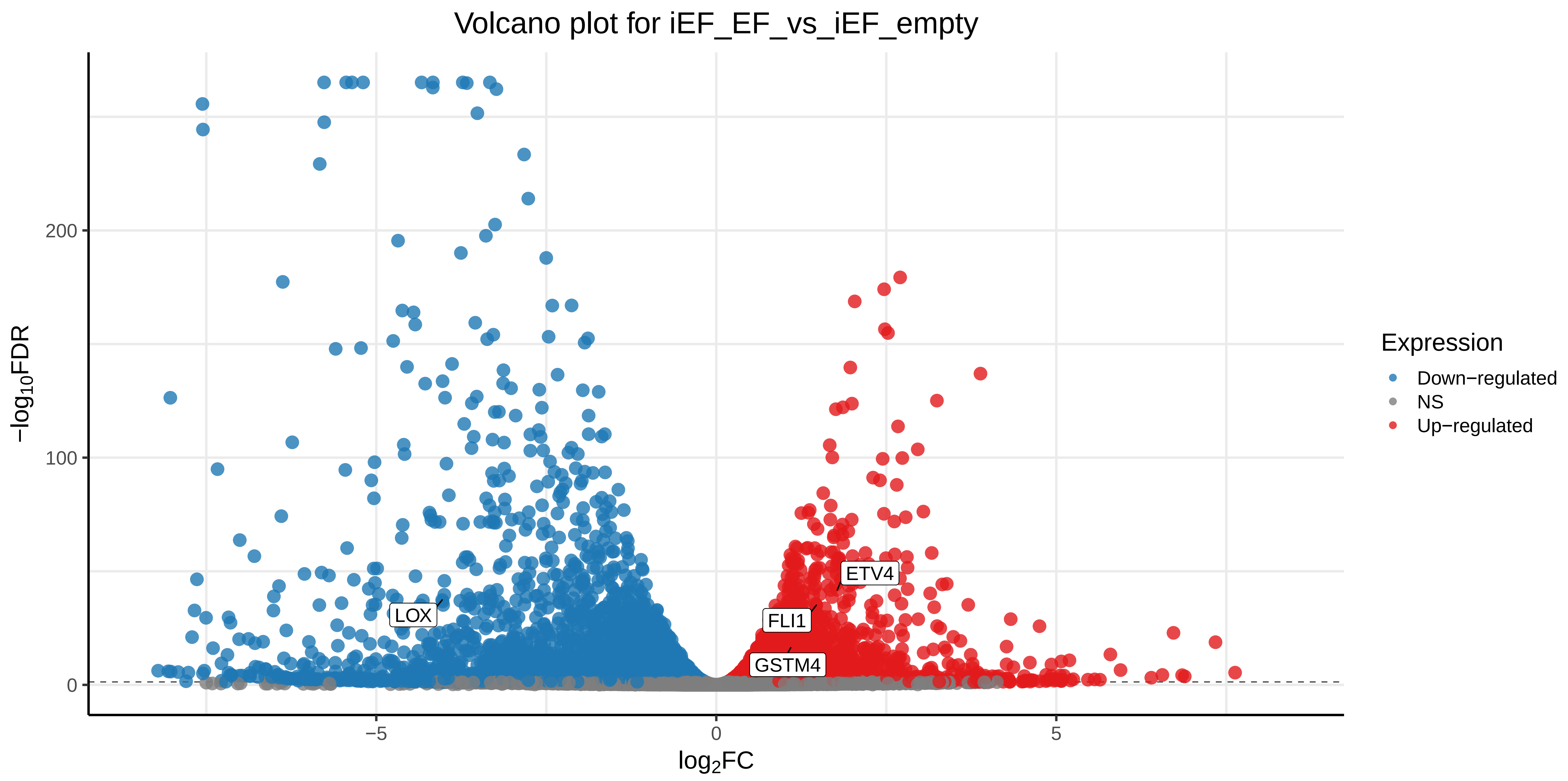

Volcano plot for iEF_EF_vs_iEF_empty
−log10FDR
200
100
0
−5
0
5
log2FC
LOX
FLI1
GSTM4
ETV4
Expression
Down−regulated
NS
Up−regulated

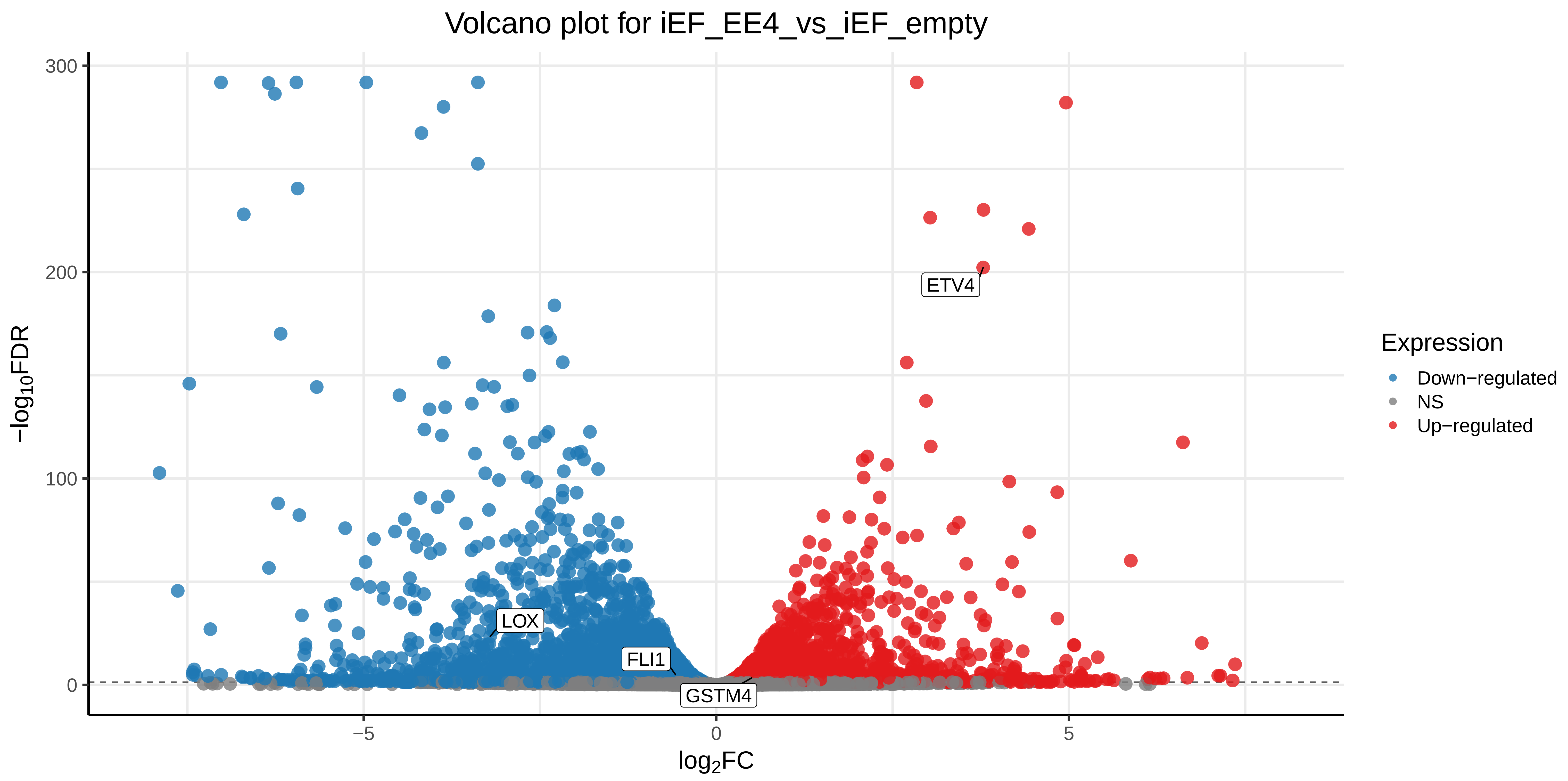

Volcano plot for iEF_EE4_vs_iEF_empty
300
200
100
0
−log10FDR
−5
0
5
log2FC
ETV4
LOX
FLI1
GSTM4
Expression
Down−regulated
NS
Up−regulated
